# Metagenomic profiling and transfer dynamics of antibiotic resistance determinants in a full-scale granular sludge wastewater treatment plant

**DOI:** 10.1101/2022.03.01.482492

**Authors:** David Calderón-Franco, Roel Sarelse, Stella Christou, Mario Pronk, Mark C.M. van Loosdrecht, Thomas Abeel, David G. Weissbrodt

**Affiliations:** Department of Biotechnology, Delft University of Technology, Delft, The Netherlands; Royal HaskoningDHV, Amersfoort, The Netherlands; Delft Bioinformatics Lab, Delft University of Technology, Delft, The Netherlands; Infectious Disease and Microbiome Program, Broad Institute of MIT and Harvard, Cambridge, USA

**Author notes:** Correspondence: David G. Weissbrodt, Assistant Professor, Weissbrodt Group for Environmental Life Science Engineering, Environmental Biotechnology Section, Department of Biotechnology, Delft University of Technology, van der Maasweg 9, 2629 HZ Delft, The Netherlands. Shared senior authorship (Thomas Abeel, David G. Weissbrodt).

**Keywords:** Aerobic granular sludge, Free-floating extracellular DNA, Intracellular DNA, Antibiotic resistance genes, Mobile Genetic Elements, qPCR, Metagenomics

## Abstract

In the One Health context, wastewater treatment plants (WWTPs) are central to safeguard water resources. Nonetheless, many questions remain about their effectiveness to prevent the dissemination of antimicrobial resistance (AMR). Most surveillance studies monitor the levels and removal of selected antibiotic resistance genes (ARGs) and mobile genetic elements (MGEs) in intracellular DNA (iDNA) extracted from WWTP influents and effluents. The role of extracellular free DNA (exDNA) in wastewater is mostly overlooked. In this study, we analyzed the transfer of ARGs and MGEs in a full-scale Nereda® reactor removing nutrients with aerobic granular sludge. We tracked the composition and fate of the iDNA and exDNA pools of influent, sludge, and effluent samples. Metagenomics was used to profile the microbiome, resistome, and mobilome signatures of iDNA and exDNA extracts. Selected ARGs and MGEs were analyzed by qPCR. From 2,840 ARGs identified, the genes *arr-3* (2%)*, tetC* (1.6%)*, sul1* (1.5%)*, oqxB* (1.2%), and *aph(3”)-Ib* (1.2%) were the most abundant among all sampling points and bioaggregates. *Pseudomonas*, *Acinetobacter*, *Aeromonas*, *Acidovorax*, *Rhodoferax,* and *Streptomyces* populations were the main hosts of ARGs in the sludge. In the effluent, 478 resistance determinants were detected, of which 89% from exDNA potentially released by cell lysis during aeration in the reactor. MGEs and multiple ARGs were co-localized on the same extracellular genetic contigs. These can pose a risk for AMR dissemination by transformation into microorganisms of receiving water bodies. Total intracellular ARGs decreased 3-42% as a result of wastewater treatment. However, the *ermB* and *sul1* genes increased by 2 and 1 log gene copies mL^-1^, respectively, in exDNA from influent to effluent. The exDNA fractions need to be considered in AMR surveillance, risk assessment, and mitigation.

**Graphical abstract:** 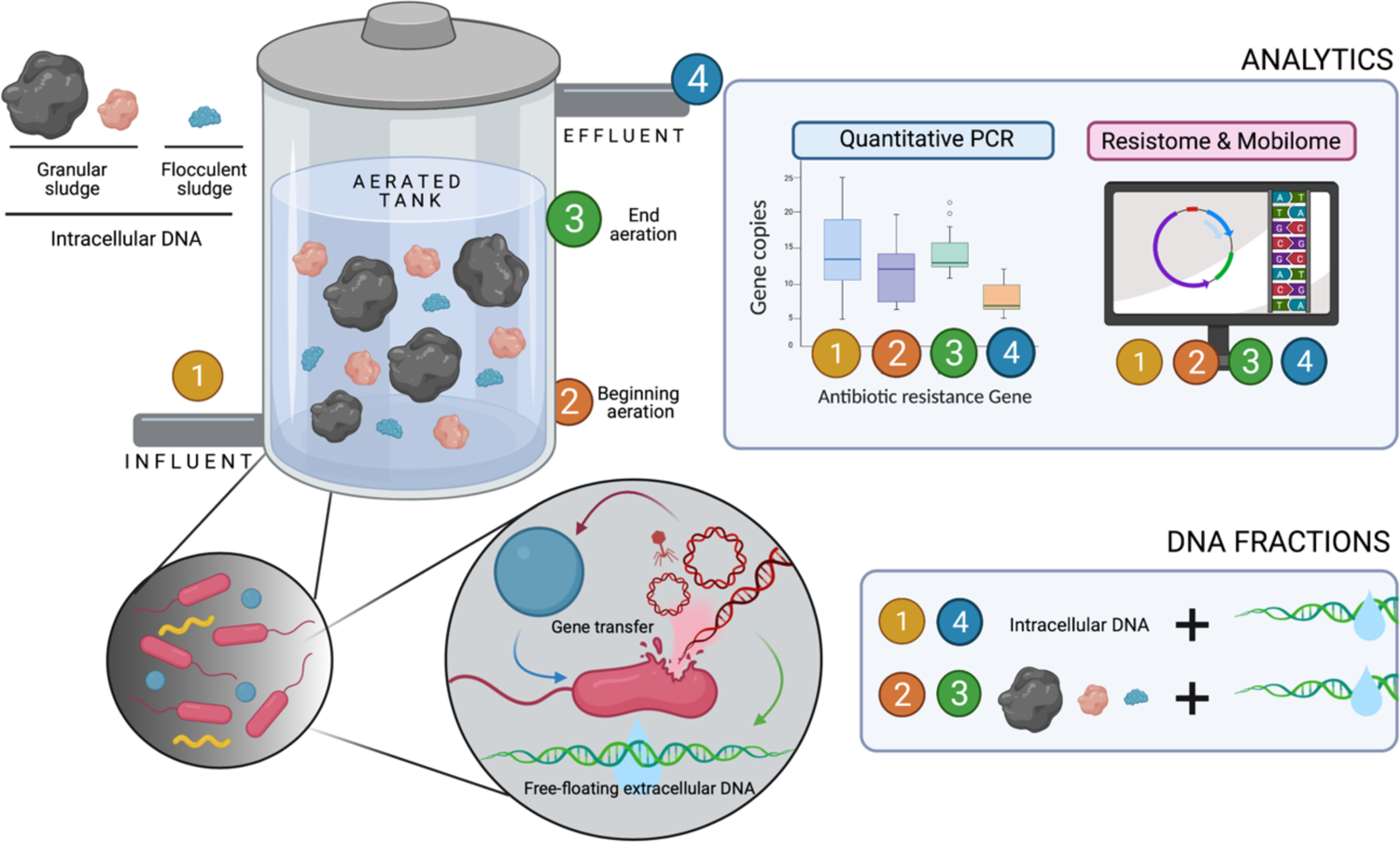

**Highlights:**

- A DNA database from an AGS reactor was constructed to study the system resistome, mobilome, and microbiome.
- The genera *Pseudomonas* and *Rhodoferax* were the predominant ARG carriers in the system.
- MGEs and ARGs often co-localize on contigs recovered from the exDNA of the effluent.
- AGS plants are efficient at reducing ARB.
- The exDNA is an underestimated DNA fraction containing ARGs in the effluent.

## 1. Introduction

Drug-resistant diseases currently cause at least 700,000 deaths globally per year. This number could increase to 10 million y^-1^ by 2050 across all income regions, under the most alarming scenario, which is that no action is taken to contain antimicrobial resistance (AMR) (WHO & UN, 2019). The proliferation of antibiotic-resistant bacteria (ARB) directly correlates with the widespread use of corresponding antibiotics (Davies and Davies, 2010). Medically problematic pathogens that acquired multidrug resistance through misuse of antibiotics are bacteria responsible for tuberculosis, acute respiratory infections, sexually transmitted infections, and bacillary dysentery (Ukuhor, 2021).

Within the One Health context, wastewater treatment plants (WWTPs) should form a barrier between sewage that transports high loads of antibiotics and antibiotic resistance genes (ARGs) emitted by anthropogenic metabolisms, and aquatic ecosystems (Bürgmann et al., 2018). Microorganisms in WWTPs biological processes are often considered reservoirs of ARGs, while also supposed to contribute in mitigating AMR dissemination by degrading antibiotics and AMR determinants. However, current WWTP designs are not optimized to this end.

Currently, AMR surveillance in wastewater is primarily conducted with molecular biology measurements that target the intracellular DNA (iDNA) of ARB. By examining the influent and effluent of 62 Dutch WWTPs by qPCR, Pallares-Vega et al. (2019) observed not only a reduction in the abundance of ARGs, but also an increase in the relative abundance of resistance plasmids of the incompatibility group 1 (IncP-1). Guo et al. (2017) used metagenomics to describe microbiome, mobilome, and resistome patterns form the iDNA pool of a Chinese WWTP, revealing that *Clostridium* and *Nitrosomonas* can carry ARGs during WWT.

Besides identifying at high resolution the hosts of ARGs in microbial communities of activated sludge, it remains primordial to elucidate the mechanisms and mobile genetic elements (MGEs) that transfer AMR determinants in these populations.

In addition to iDNA, extracellular free DNA (exDNA) contains a high proportion of MGEs (Calderón-Franco et al., 2020). Different genetic structures and architectures (plasmids, transposons, insertion sequences, and integrases, among others) transfer ARGs between bacteria, but many questions remain. The analysis of both intracellular ARGs (iARGs) and extracellular ARGs (exARGs) combined with mobilome co-localization analysis has rarely been performed in complex environmental samples such as wastewater biomasses. Such co-localization analysis involves the characterization of the resistome fraction in genomic proximity to horizontal gene transfer (HGT) mediators such as plasmids and other mobile genetic elements (Slizovskiy et al., 2020).

Apart from the presence of exDNA in wastewater, not much is known about the actual transfer of AMR in the microbiome of WWTPs and the underlying effects of biofilms. Dense microbial aggregations can promote the horizontal transfer of antibiotic resistance genes (ARGs) in aquatic environments (Abe et al., 2020; Madsen et al., 2012). Full-scale Nereda® plants that use aerobic granular sludge (AGS) for an integral removal of nutrients (Pronk et al., 2015) harbor a hybrid sludge composed of granules (0.2-3 mm) that have similar properties as biofilms. Therefore, AGS SBRs form interesting microbial ecosystems to study the fate of AMR. Metagenomic analysis during granulation at pilot scale showed that ARGs enriched in both iDNA and exDNA fractions of AGS during the granule development stage, and that integrons played an essential role in carrying exARGs (Li et al., 2020).

In this study, we analyzed the transfer dynamics of ARGs and MGEs in a full-scale AGS Nereda® plant. Metagenomic and qPCR analyses were performed on the iDNA and exDNA pools of samples collected from the influent, different sludge fractions and the effluent. The sludge was sampled over the different SBR phases (end of anaerobic feeding, end of aeration) and sifted for different sized bioaggregates (flocs, small and big granules), to track transfer phenomena. The findings highlight the fate of the AMR determinants in a full-scale AGS WWTP and the importance of considering both iDNA and exDNA pools in AMR dissemination studies.

## 2. Materials and methods

### 2.1. Sampling of a full-scale AGS WWTP

The sampling campaign performed on a Nereda® plant located in Utrecht (52.11215, 5.10813), The Netherlands. Sampling was performed over 3 days (Monday, Wednesday, and Friday) within the same week, under dry weather conditions without significant variations in hydraulic retention time (20 hours). Since the sludge retention time was relatively long (30 days in average; >40 days for large granules and <10 days for small granules and flocs) in this installation, the 3 sampling days were considered as biological replicates.

Biological samples were collected every day from pre-settled influent wastewater, from the mixed liquor at the beginning (*i.e.*, after anaerobic feeding of the wastewater) and end of the aeration phase, and from the effluent of the SBR (**Figure 1a**). Total volumes of 1 L of influent and 1 L of effluent were collected as a 24-h flow-proportional composite samples. For the other process points, the sludge was sampled from the top of the tank as grab samples of 1 L with a pole container during the daily operations of the SBR. All samples were transported in a cooled container and processed within a time frame of less than 4 h prior to DNA extraction.

**Figure 1.**
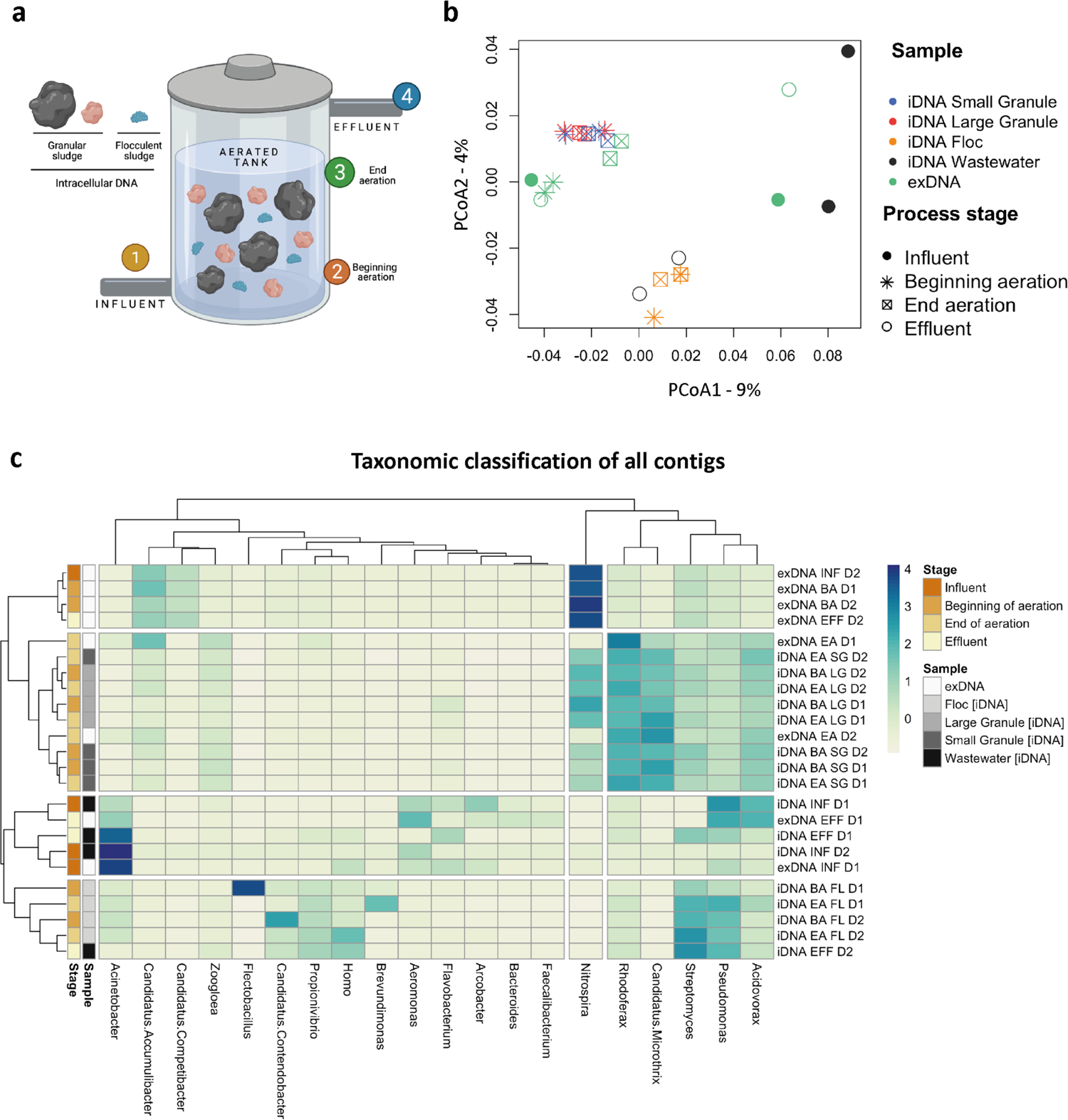
**(a)** Schematic representation of an AGS SBR, highlighting biomass metrics and sampling points 1-4. **(b)** PCoA of the absolute microbiome composition at genus-level after normalization and variance stabilization transformation, with Bray-Curtis as distance metric. Colors indicate microbial fractions: iDNA small granule (blue), iDNA large granule (red), iDNA floc (orange), iDNA wastewater (black, including influent and effluent samples), and exDNA (green). Symbols indicate stages in WWT operation: Influent (filled circles), beginning of aeration (asterisks), end of aeration (crossed squares), and effluent (empty circles). **(c)** Taxonomic classification at genus-level heatmap. Colors represent the Z-score computed from the relative abundance at genus-level present above 3% in all classified sequences.

The granular sludge samples were sieved in 3 different fractions of bioaggregates (flocs <0.2 mm, small granules of 0.2-1 mm, and large granules >1 mm) according to Ali et al. (2019). The water at the outlet of the sieving unit was used for exDNA isolation. The fractions of exDNA were isolated in a time-frame of less than 4 h after sampling. The sieved fractions of granular sludge were stored at −20°C in a time-frame of less than 4 h after pending isolations of iDNA.

In total, 12 analytes were obtained and sequenced by metagenomics per sampling day, *i.e.*, 36 analytes for the whole campaign. **Table 1** summarizes the set of samples and analytes.

**Table 1.**
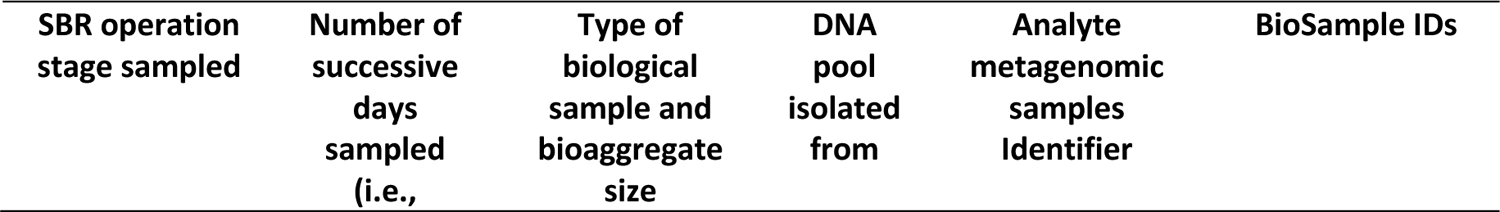
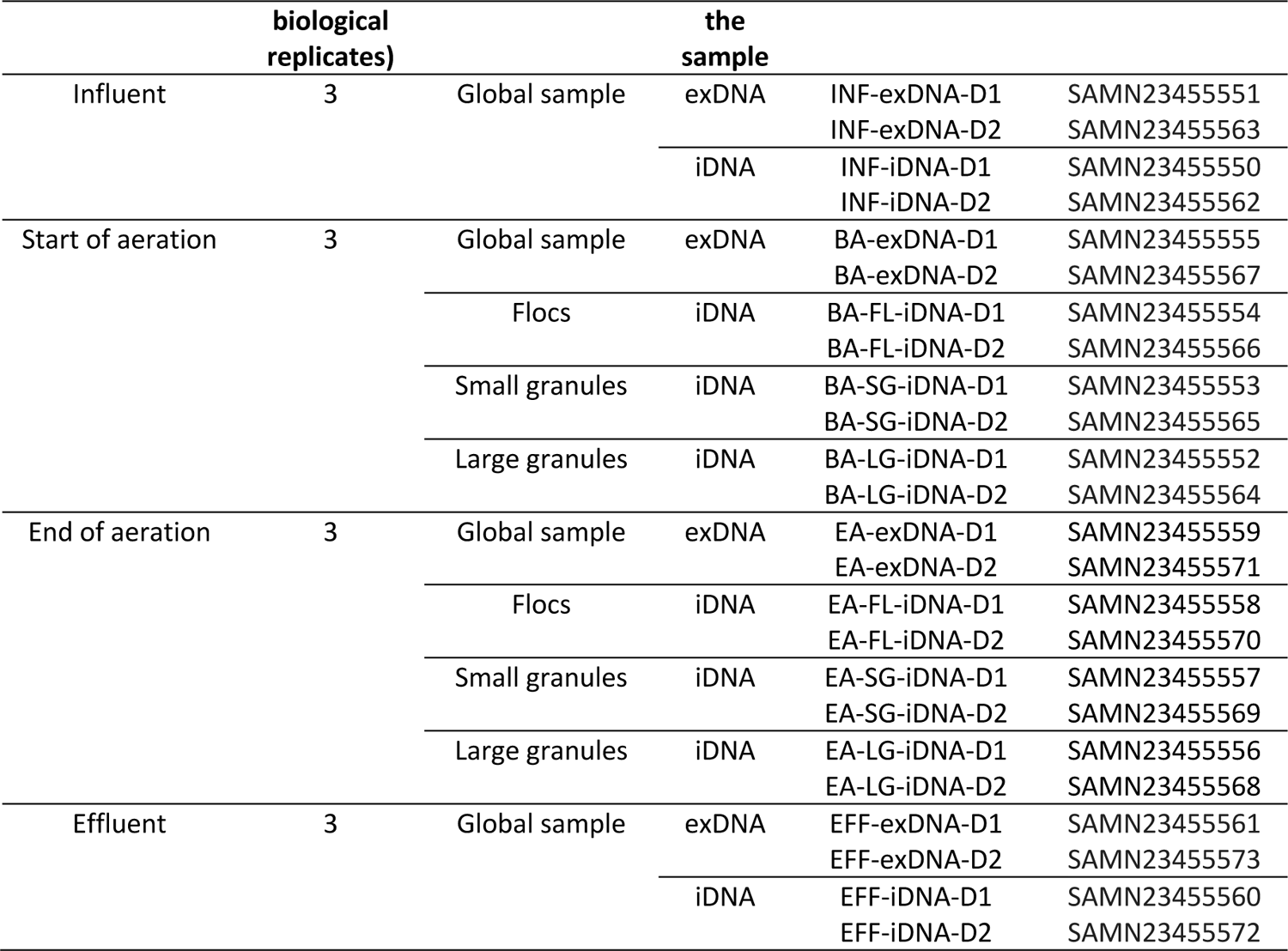
Samples collected from the influent, sludge, and effluent of the SBR during each of the 3 sampling days at WWTP Utrecht. Granular sludge samples were sieved in 3 different fractions of flocs, small granules, and large granules. The pools of intracellular DNA (iDNA) and free-floating extracellular DNA (exDNA) extracted from each sample as analytes for metagenomics are indicated. Two out of the three biological replicates were sequenced for metagenomics analysis, indicated as “-D1” and “-D2” for samples taken on Monday and Friday, respectively.

| SBR operation stage sampled | Number of successive days sampled (*i.e.*, | Type of biological sample and bioaggregate size | DNA pool isolated from | Analyte metagenomic samples Identifier | BioSample IDs |
| --- | --- | --- | --- | --- | --- |

|  | biological replicates) |  | the sample |  |  |
| --- | --- | --- | --- | --- | --- |
| Influent | 3 | Global sample | exDNA | INF-exDNA-D1<br>INF-exDNA-D2 | SAMN23455551<br>SAMN23455563 |
|  |  |  | iDNA | INF-iDNA-D1<br>INF-iDNA-D2 | SAMN23455550<br>SAMN23455562 |
| Start of aeration | 3 | Global sample | exDNA | BA-exDNA-D1<br>BA-exDNA-D2 | SAMN23455555<br>SAMN23455567 |
|  |  | Flocs | iDNA | BA-FL-iDNA-D1<br>BA-FL-iDNA-D2 | SAMN23455554<br>SAMN23455566 |
|  |  | Small granules | iDNA | BA-SG-iDNA-D1<br>BA-SG-iDNA-D2 | SAMN23455553<br>SAMN23455565 |
|  |  | Large granules | iDNA | BA-LG-iDNA-D1<br>BA-LG-iDNA-D2 | SAMN23455552<br>SAMN23455564 |
| End of aeration | 3 | Global sample | exDNA | EA-exDNA-D1<br>EA-exDNA-D2 | SAMN23455559<br>SAMN23455571 |
|  |  | Flocs | iDNA | EA-FL-iDNA-D1<br>EA-FL-iDNA-D2 | SAMN23455558<br>SAMN23455570 |
|  |  | Small granules | iDNA | EA-SG-iDNA-D1<br>EA-SG-iDNA-D2 | SAMN23455557<br>SAMN23455569 |
|  |  | Large granules | iDNA | EA-LG-iDNA-D1<br>EA-LG-iDNA-D2 | SAMN23455556<br>SAMN23455568 |
| Effluent | 3 | Global sample | exDNA | EFF-exDNA-D1<br>EFF-exDNA-D2 | SAMN23455561<br>SAMN23455573 |
|  |  |  | iDNA | EFF-iDNA-D1<br>EFF-iDNA-D2 | SAMN23455560<br>SAMN23455572 |

### 2.2. Extractions of exDNA and iDNA analytes

The pools of iDNA and exDNA were differentially isolated for all samples, as recently described by Calderón-Franco et al. (2020).

#### Filtrations of influents and effluent samples

The total volumes of 1 L of influent and effluent were filtered through 0.22 μm polyvinylidene fluoride (PVDF) membranes (Pall Corporation, USA). The filtrates were used to isolate the exDNA. The biomasses retained on membranes were used for iDNA extractions.

#### Centrifugations and homogenizations of sludge fractions

The sieved sludge fractions were centrifuged at 4,000 ×g for 10 min before storing at −20°C. The biomass pellets were used for iDNA extraction. The pellets were first treated with DNAse I to remove remaining exDNA fragments, and homogenized with a Potter-Elvehjem pestle (Wheaton, USA).

#### Isolations of exDNA

The water fractions remaining after sieving were used for exDNA isolation. After filtration through 0.22 µm PVDF membranes, exDNA was loaded for ion-exchange chromatography on a 1-mL, positively charged, diethylaminoethyl cellulose (DEAE) column with 2-µm pore size (BIA Separations, Slovenia). Manufacturer’s instructions were followed to condition the column and to process the chromatographic operations. The following buffers and solutions were used to equilibrate, elute, regenerate, clean, and store the DEAE column: (i) equilibration buffer (50 mmol L^−1^ Tris, 10 mmol L^−1^ EDTA at pH 7.2); (ii) elution buffer (50 mmol L^−1^ Tris, 10 mmol L^−1^ EDTA, 1.5 mol L^−1^ NaCl at pH 7.2); (iii) regeneration buffer (50 mmol L^−1^ Tris, 10 mmol L^−1^ EDTA, 2 mol L^−1^ NaCl at pH 7.2); (iv) cleaning solution (1 mol L^−1^ NaOH and 2 mol L^−1^ NaCl); (v) storage solution (20% ethanol in ultrapure water). The column was equilibrated using the equilibration buffer at a flowrate of 0.6 mL min^−1^ to maintain the pressure below the maximum limit of 1.8 MPa. After equilibration, the entire volume (1 L) of filtrates containing the exDNA was loaded in the DEAE column using an HPLC pump (Shimadzu Corporation, Japan) at the same flowrate of 0.6 mL min^−1^. After retention, exDNA was eluted at a flow rate of 1 mL min^−1^ using the elution buffer and tracked over time with an HPLC UV-Vis photodiode array detector (Waters Corporation, USA) recording the UV-light absorption of DNA at 260 nm. The recovered raw exDNA was precipitated from the eluate with ethanol (Moore and Dowhan, 2002). The residual proteins bound to exDNA were digested by 2-h incubation with 0.85 g L^−1^ proteinase K (Sigma-Aldrich, UK) at room temperature. This enzymatic reaction was stopped at 50°C for 10 min in a thermoblock (ThermoMixer, Eppendorf). The protein-digested exDNA was finally purified using GeneJET NGS Cleanup Kit (Thermo Scientific, USA). The purified exDNA isolates were stored at −20°C pending analysis.

#### Extractions of iDNA

The membranes containing the biomasses filtrated from the influent and effluent samples were directly used for iDNA extraction using PowerWater DNA extraction kit (QIAGEN, The Netherlands) following manufacturer’s instructions. For the bioaggregate samples, a mass of 0.25 g of wet weight of homogenized biomass was used to extract iDNA. IDNA extractions were performed using the DNeasy PowerSoil DNA extraction kit (QIAGEN, The Netherlands) following manufacturer’s instructions.

### 2.3. Library preparation, sequencing, quality control, assembly, and binning

#### Selection of analytes

From the set of 36 analytes of exDNA and iDNA obtained over the 3 days, the analytes from the Monday and Friday (*i.e.*, 24 analytes) were selected as duplicates (economic optimum) for metagenomics. These exDNA and iDNA analytes were sent to DNASense (Aalborg, Denmark) for library preparation and sequencing.

#### Preparation of libraries

The DNA was quantified using Qubit (Thermo Fisher Scientific, USA) and fragmented to approximately 550 bp using a Covaris M220 with microTUBE AFA Fiber screw tubes and the settings: Duty Factor 10%, peak/displayed power 75 W, cycles/burst 200, duration 40 s and temperature 20°C. The fragmented DNA was used for metagenome preparation using the NEB Next Ultra II DNA library preparation kit.

#### Sequencing of libraries

Libraries were sequenced using the HiSeq sequencer (Illumina, USA) as 2×150 bp paired-end reads.

#### Quality control of sequence reads

After sequencing, a dataset containing 48 paired-end read samples with an average of 16 million reads per sample was obtained. Metagenomics workflow is summarized in **Figure S1** in supplementary information. The minimum and maximum numbers of quality-filtered, non-duplicated sequencing reads of 150 bp ranged from 12 to 18 million (**Figure S2** in supplementary information). The quality of the Illumina reads was assessed using FastQC version 0.11.9 with default parameters (Andrews, 2010). Low-quality paired-ends reads were trimmed and filtered by Trimmomatic version 0.39 on paired-end mode (Bolger et al., 2014).

#### Assembly of sequence reads

The trimmed reads were assembled into contigs using metaSPAdes version 3.14.1 (Nurk et al., 2017) on meta mode on default parameters.

#### Binning of DNA contigs

Contigs resulting from the sequencing of only the iDNA pools of bioaggregates were binned with MetaBAT version 2.2.15 (Kang et al., 2019) to reconstruct metagenome-assembled genomes (MAGs) on default parameters. The MAGs were used to analyze secondary metabolite biosynthesis gene clusters.

### 2.4. Generation of the taxonomic database of aerated granular sludge

A database of sequences from MAGs, contigs, and reads specific to the microbial environment of AGS was built to accurately profile the microbiome. The Kraken 2.0 standard database of 9.1 Gbp of genomic sequences (Wood et al., 2019) was used as a basis. However, because complete genomes of the organisms found in AGS systems were often unavailable in the Kraken 2.0 database, additional genetic fragments specific to some taxa were added to the database. As an example, abundant genera in AGS like *Tetrasphaera*, *Trichococcus* and “*Candidatus* Accumulibacter” were not or poorly annotated in the standard database used in Kraken2.0. In total, 94,005 sequences of 2,223 unique taxa were added to the Kraken 2.0 standard database.

The taxa of the AGS samples were classified from both short reads and contig sequences by combining the MetaPhlAn3.0 (Truong et al., 2015), MG-RAST (Meyer et al., 2018) and Kraken2.0 (Wood et al., 2019) computational tools for phylogenetic analysis of metagenomics data with their corresponding databases, as well as BLASTn to align contigs of >1500 bp against the MiDAS database of 16S rRNA gene sequences with a cut-off e-value <10^-5^ and sequence identity >97% [https://www.midasfieldguide.org/] (Nierychlo et al., 2020). A literature study was conducted in which taxa were added to the database if theoretically present above 1% relative abundance but not present in the database. Kraken2.0 takes all the sequences added in a selected database and finds k-mers (short genomic substrings specific for certain taxa) for further taxonomic classification.

The reads (full and partial genomes) of the predominant taxa were added to the adapted taxonomic database of AGS, when these lineages were detected above 1% of relative abundance from the metagenomics datasets using one of the previously described classification tool but not present in the Kraken2.0 database. This was performed on taxonomic levels from phylum to species.

### 2.5. Microbiome profiling of iDNA and exDNA pools obtained from the AGS process

Kraken2.0 with short-reads as input was selected to profile the microbiome. Classification with Kraken2.0 was performed on pair-end mode on the quality-controlled short reads, using the newly constructed database.

The taxonomic classification was also performed on contigs >500 bp that were identified to contain ARGs (see § 2.6 hereafter), in order to determine potential ARGs hosts.

The taxonomic classification outcomes from Kraken2.0 were converted into abundance tables using the Pavian visualization tool (Breitwieser and Salzberg, 2020) to explore metagenomics classification datasets. Heatmaps were generated with the R package “pheatmap” (Kolde, 2019).

### 2.6. Resistome and mobilome profiling of iDNA and exDNA obtained from the AGS process

ARGs were annotated by aligning the assembled contigs >500 bp to the ResFinder 4.0 resistance gene database using the BLASTn nucleotide alignment tool with a cut-off E-value of <10^-5^ and sequence identity above 90% (Bortolaia et al., 2020).

MGEs were classified on the same set of contigs >500 bp using BLASTn by aligning them with the same cut-off and sequence identity setpoints to the following databases depending to the types of MGEs. Bacterial insertion sequences were identified by aligning against the ISfinder database (Siguier et al., 2006). Integrons, integrases, and gene cassettes were identified using the INTEGRALL database (Moura et al., 2009). Bacterial integrative and conjugative elements were identified using the ICEBERG database (Liu et al., 2019).

For all queries, the ARG or MGE identified with the best score (*i.e.*, equal to the sequence identity multiplied by the coverage factor) was selected to annotate the query.

BLASTn was performed with different databases (ResFinder 4.0 for ARGs and ISfinder, INTEGRALL and ICEBERG for MGEs) on the same set of contigs to identify where ARGs and MGEs co-localized. BLASTn was performed with a cut-off E-value of <10^-5^ and sequence identity above 90%. Contigs >500 bp that simultaneously contained hits from the ResFinder 4.0 database and at least one of the different MGE databases were considered to have co-localized.

The contigs that contained both ARGs and MGEs were used as input queries against the NCBI plasmid database in order to know if such contig belonged to a plasmid. A contig was identified as part of a potential plasmid if exclusively aligning with plasmids in the entire NCBI plasmid database with sequence identities >90%.

### 2.7. Functional genetic analysis of iDNA and exDNA pools obtained from the sludge

Prokka version 1.14.5 was used to annotate the assembled contigs >500 bp, with the default databases and parameters on metagenomic mode (Seemann, 2014). K-numbers were assigned to all predicted coding sequences (CDSs) using GhostKoala, after which the KEGG database was used for analyse the functional and AMR pathways (Kanehisa et al., 2002; Seemann, 2014). The antibiotics and secondary metabolites analysis shell (antiSMASH, v5.0) tool was used to identify, annotate, and analyze the secondary metabolite biosynthesis gene clusters, such as involved on antibiotics production, in MAGs binned from the iDNA bioaggregate samples (Blin et al., 2021).

### 2.8. Multidimensional scaling analysis to cluster microbiome and resistome datasets

To identify patterns between the microbiome and resistome, different types of ordination and statistical numerical methods were performed using the R software. The input data was the output of the taxonomic classification from Kraken2.0 with the adapted database.

Dimension reduction is a projection-based method that transforms the data by projecting it onto a set of perpendicular axes. Before dimension reduction, the metagenomic datasets were normalized by dividing the measurements by the corresponding sample size factor using software package “edgeR” (Robinson et al., 2009), and a variance stabilization transformation (VST) was performed using “DESeq2” (Love et al., 2014), respectively. The “vegan” package (Dixon, 2003) transformed the stabilized matrices into distance matrices. The Bray-Curtis distance metric was identified as the optimal distance metric for all dimension reduction methods.

The dimensions of the metagenomics datasets were reduced using Non-metric Multidimensional scaling (NMDS), Principal Coordinates Analysis (PCoA), and t-distributed stochastic neighbor embedding (t-SNE). These methods were applied to the taxonomic data at phylum and genus levels, and to the AMR group data. For all methods, scree-plots, stress-plot, and Shepard-plots were used to visualized the data accurately in two dimensions. t-SNE was applied using the “tsne” package, and a maximum of 2,000 iterations and a perplexity parameter of 5 as input (van der Maaten and Hinton, 2008). PCoA, t-SNE, and NMDS were performed on different taxonomic levels and on the resistome data (**Figure S3**). All dimension reductions yielded similar clustering patterns, giving equivalent results for each sampling point and sample type. PCoA was therefore chosen as representative analysis to characterize the similarities and differences between samples.

### 2.9. Quantitative PCR of selected ARGs

To evaluate the WWTP performance in terms of removing ARGs and MGEs, to examine their presence during the different process steps, and to track their transfer between iDNA and exDNA fractions, a molecular analysis by quantitative polymerase chain reaction (qPCR) was applied to detect a selected panel of genes (**Table 2**) from the three biological replicates (Monday, Wednesday, and Friday) of the samples listed in **Table 1**.

**Table 2.** ARGs and MGE selected for qPCR with their description

| Category | Gene | Description |
| --- | --- | --- |
| Bacteria normalization | *16S rRNA* | RNA component of the 30S small subunit of prokaryotic ribosome |
| | *rpoB* | Bacterial RNA polymerase subunit $\beta$ |
| ARGs | *bla<sub>CTXM</sub>* | Bla-cefotaxime-hydrolyzing β-lactamase |
|  | *ermB* | Erythromycin resistance via methylation of 23S rRNA |
|  | *qnrS* | Quinolone resistance protein |
|  | *sul1* | Sulfonamide resistant dihydropteroate synthase |
|  | *sul2* | Sulfonamide resistant dihydropteroate synthase |
|  | *tetO* | Tetracycline resistance protein TetO |
| MGE | *Int1* | Class 1 integron integrase |

All iDNA and exDNA analytes isolated from the three-biological sample from the influent, beginning and end of aeration, and effluent of the SBR were used for qPCR. Each analyte was measured in technical duplicates by qPCR. The *16s rRNA* and *rpoB* reference genes were quantified for the normalization of gene copies to the concentration of bacteria in the iDNA samples.

Based on antibiotics consumption data in the Netherlands (RIVM and SWAB, 2020), ARGs were targeted from the antibiotic groups of beta-lactams (*bla_CTXM_*), macrolides (*ermB*), fluoroquinolones (*qnrs*), sulfonamides (*sul1/sul2*), and tetracyclines (*tetO*). The *intI1* gene encoding the class I integron-integrase was quantified. These integrase class I cassettes are related to ARG mobility, acquisition, and exchange between microorganisms (Ma et al., 2017).

Primers, thermal cycler conditions, and gBlocks gene fragments used for standards generation and quantification during qPCR are given in **Tables S1, S2,** and **S3**, respectively.

### 2.10. Analysis of qPCR results

R version 3.6.3 (R Foundation for Statistical Computing., 2018) was used for visualization and statistical analysis of qPCR data.

A non-parametric Wilcoxon test was performed to determine the significance of the differences detected in the concentrations of genes (log_10_ gene copies VSS^-1^ for iDNA or mL^-1^ for exDNA) between the iDNA and exDNA fractions, and in the absolute and relative abundances of these genes in the influent and effluent (*i.e.*, indicating a removal of the genes due to treatment).

The same Wilcoxon test was used to indicate the significant difference between *rpoB* and *16S rRNA* gene copies for their use as normalization factors in the qPCR results. Statistical significance was established at the 95% confidence level (p<0.05). The p-values for the Wilcoxon test are given in Supplementary Material (**Table S4-S5**).

## 3. Results and discussion

The iDNA and exDNA pools were isolated from samples collected over 3 sampling days (Monday, Wednesday, Friday) out of the influent, mixed liquor (flocs, small and large granules), and effluent of a full-scale AGS SBR of WWTP Utrecht. The Monday and Friday iDNA and exDNA samples were metagenomically sequenced as biological duplicates to analyze their microbiome, resistome, and mobilome. The construction of the taxonomic database of AGS environment from the metagenomes was an essential step and first result to this end. Numerical analyses were used to map and identify patterns in these metagenomic data, with a focus on the ARG and MGE in microbiome of the WWTP. All samples from the 3 days were subsequently measured by qPCR to track selected ARGs and MGEs in the different samples.

### 3.1. Taxonomic composition of exDNA mainly linked to the granular sludge fraction

After taxonomic classification at genus level of the DNA contigs assembled from the metagenomic sequencing reads, principal coordinates analysis (PCoA) and other multidimensional scaling methods (**Figure S3** in supplementary material) were efficient to observe the clustering effects in data related to the specific sample DNA fraction (**Figure 1b**).

The influent samples showed a large difference in composition for both the iDNA and exDNA pools. Microbial compositions of sewage can vary over an active week, based on the variety of emission sources and streams in a wastewater catchment area and environmental factors (Pallares-Vega et al., 2021).

In the effluent, a lower variability between the two samples was observed for the iDNA fraction. The taxonomic composition of the iDNA of the effluent highly resembled the composition of the iDNA of activated sludge floc fraction in the treatment reactor. On the other hand, the phylogenetic signatures of the exDNA fraction of the effluent showed a larger variation between the samples.

In the wastewater treatment plant microbial cells can lyse, releasing iDNA as exDNA. The exDNA fractions might not persist as “free-floating” during the whole process. It can adhere to particles, get degraded, or taken up via natural transformation in competent cells (Dong et al., 2019; Zhou et al., 2019).

From the different size fractions of bioaggregates in the sludge, the microbial community compositions of the small (blue in **Figure 1b**) and large (red) granules were similar, while the flocs (orange) clustered separately. This is similar as the observations by Ali et al (2019) that the floc fraction is more diverse and contains a large number of bacteria originating from the influent wastewater. Microbial niches establish along gradients of substrates and other dissolved components like oxygen and nitrogen oxides inside bioaggregates (Winkler et al., 2018).

The taxonomic affiliations of the exDNA sequences of the mixed liquor sampled at the end of aeration were similar to those of the iDNA extracted from small and large granules, rather than the flocs. This suggests that exDNA was released from the granules by cell lysis during aeration. Sengar et al. (2018) and Toh et al. (2003) have shown that dead biomass releases its genetic material into the extracellular environment during the AGS process. In activated sludge, Yuan et al. (2019) have identified that the taxa reflected by the iDNA and exDNA fractions are similar, suggesting a correlation between changes in the microbial composition of the activated sludge and in exDNA. Using the same exDNA extraction methodology, we have shown in a previous study that exDNA present in activated sludge mainly originates from decaying microbial populations rather than from transport from the influent (Calderón-Franco et al., 2020).

### 3.2. AGS microbiome and involvement in AMR dissemination

The patterns of relative abundances of genera in the different stages of the AGS process and in the iDNA and exDNA fractions were examined by constructing a heatmap (**Figure 1c**) for the clusters observed by PCoA.

Using the standard Kraken2.0 database (NCBI based), 2,548 genera were identified from the whole set of iDNA and exDNA analytes: 57 genera were more abundant than 1% of sequencing read counts, and 15 of those were predominant above 3% in at least one of the samples (**Figure S4, * symbols**). The average annotation rate of the reads was 24.7 ± 3.2 %, *i.e.*, leaving three quarters of the taxonomic information of the reads as to uncover.

To increase the reads annotation rate, a new database of genetic sequences of the AGS microbial environment was constructed. With this database, the average annotation rate was 27.1 ± 3.4 %; 2,679 genera were identified: 55 genera were above 1% relative abundance and 21 were above 3% in at least one of the analytes (**Figure S4**, ◊). Although this new annotation rate was only 3% higher than the standard database, important AGS genera were now identified. Metagenomic studies can be hindered by low annotation rates as a consequence of the high fraction of natural microbes that have not been included in the databases. The current development of databases of high-quality MAGs and properly curated databases will help enhance annotation rates in the future (Singleton et al., 2021).

From **Figure 1b**, the exDNA taxonomic compositions at the beginning of aeration resembled the exDNA from influent while at the end of aeration from both days closely resembled the iDNA fractions from small and large granules. Flocs displayed a unique profile. While the phylogenic distribution of the effluent exDNA resembled the granule phylogeny, the effluent iDNA resembled the flocs (from both beginning and end of aeration). During the fill/draw operation of the SBR, flocs leave the tank with the selection spill, therefore reducing their retention time in the process (Guo et al., 2020).

The diversity of abundant populations in the AGS detected by metagenomics is listed in **Figure 1c**. Several of these populations are involved in the conversions of C-N-P nutrients in the microbial ecosystem of AGS (Weissbrodt et al., 2014; Winkler et al., 2018), like “*Ca.* Accumulibacter”, *Tetrasphaera*, *Dechloromonas* (C, P), “*Ca*. Competibacter”, *Zoogloea, Xanthomonas*, *Rhodoferax*, *Pseudomonas, Acidovorax*, *Comamonas*, *Rhizobium* (C, N), *Nitrospira* (N), and *Acinetobacter* (C) among others. Some relate to filamentous bulking phenomena like “*Ca.* Microthrix”, *Bulkholderia*, *Nocardioides*, Kouleothrix. Some thrive on metabolites and lysis products from other organisms like *Flavobacterium*, *Bacteroides*, and *Hydrogenophaga* among many others. The metabolic functions of other organisms in the AGS remains to be uncovered. Less beneficial organisms are *Arcobacter* and *Aeromonas* that are known pathogens. *Arcobacter* crosses WWTPs from influent to effluent without settling properly (Kristensen et al., 2020). The presence in the influent of exDNA affiliating with “*Ca.* Accumulibacter” can relate to the recirculation line coming from the sludge thickener.

Among the different populations identified from the AGS (**Figure 1c**), several are known to carry AMR-related genes. *Acidovorax* (3.5 ± 0.6%) is a known carrier of beta-lactamase resistance plasmids in activated sludge samples (Zhang et al., 2011). *Rhodoferax* (3.5 ± 0.5%) has been identified to be of high relative abundance in wastewater effluent, while often identified as a carrier of resistance genes (Zhou et al., 2020). *Nitrospirae* (4.1% ± 1.1%) may play a role in AMR dissemination because it is a known host of iARGs and exARGs in WWTPs (Zhou et al., 2019). Guo et al. (2017) used metagenomics to identify *“Candidatus* Accumulibacter*”* (2.1 ± 0.5%) as a possible carrier of resistance genes over the WWT operation. *Aeromonas* was relatively abundant in all our samples (1.88 ± 0.84%). Notably, the three most critical antibiotic-resistant pathogens as designated by the World Health Organization are

*Acinetobacter baumannii, Pseudomonas aeruginosa,* and *Enterobacteriaceae* (Tacconelli and Magrini, 2017). *Acinetobacter* and *Pseudomonas* were abundant across samples (**Figure 1c** and **Figure S4**). Recovering and annotating MAGs for these populations out of AGS biomass could help identify whether they harbour pathogenic traits or not.

Interestingly, the relative abundances of exDNA sequences affiliating with the genera “*Candidatus* Microthrix” (exDNA at beginning of aeration 0.5 ± 0.1% *vs*. at end of aeration 5.6 ± 2.0%), *Acidovorax* (1.2 ± 0.1% *vs.* 4.3 ± 0.2%), *Rhodoferax* (1.8 ± 0.2% *vs.* 7.9 ± 1.3%), and *Zoogloea* (0.8 ± 0.2% *vs*. 2.6 ± 0.5%) increased over the aeration period. These filamentous bacteria (“*Ca.* Microthrix”) and denitrifiers were abundant >3% in the granule fractions of the SBR but not in flocs. These microbial populations are known to populate AGS used for full biological nutrient removal (Winkler et al., 2018).

The presence of exDNA in the water phase can relate to cell lysis or to “active” release from the cells. *Acinetobacter* (iDNA 6.1 ± 2.8%; exDNA 3.4 ± 2.2%)*, Flavobacterium* (1.2 ± 0.4%; 1.0 ± 0.5%), and *Pseudomonas* (4.3 ± 0.9%; 4.0 ± 1.1%) secrete DNA in the extracellular environment during growth in liquid media (Pietramellara et al., 2009).

Further mechanistic insights, using well-controlled experiments with systems microbiology methods, are required to identify whether cell decay or active secretion can explain the release of exDNA from microorganisms of the sludge.

### 3.3. A wide range of ARGs are present in the effluent exDNA

The resistome of wastewater systems exhibits a large diversity of ARGs. As many as 2,840 ARGs were identified from all the samples, belonging to 15 antibiotic subgroups. ARGs affiliating with MLS (Macrolide, Lincosamide, and Streptogramin; n = 910 reads), aminoglycosides (n = 598 reads), sulphonamides (n = 375 reads), beta-lactam (n = 330 reads), and tetracycline (n = 233 reads) were the most abundant (**Figure 2a**). The resistome profiles matched with Hendriksen et al. (2019): macrolides are the most abundant ARGs in urban sewage in Europe, followed by aminoglycosides and beta-lactams.

**Figure 2.**
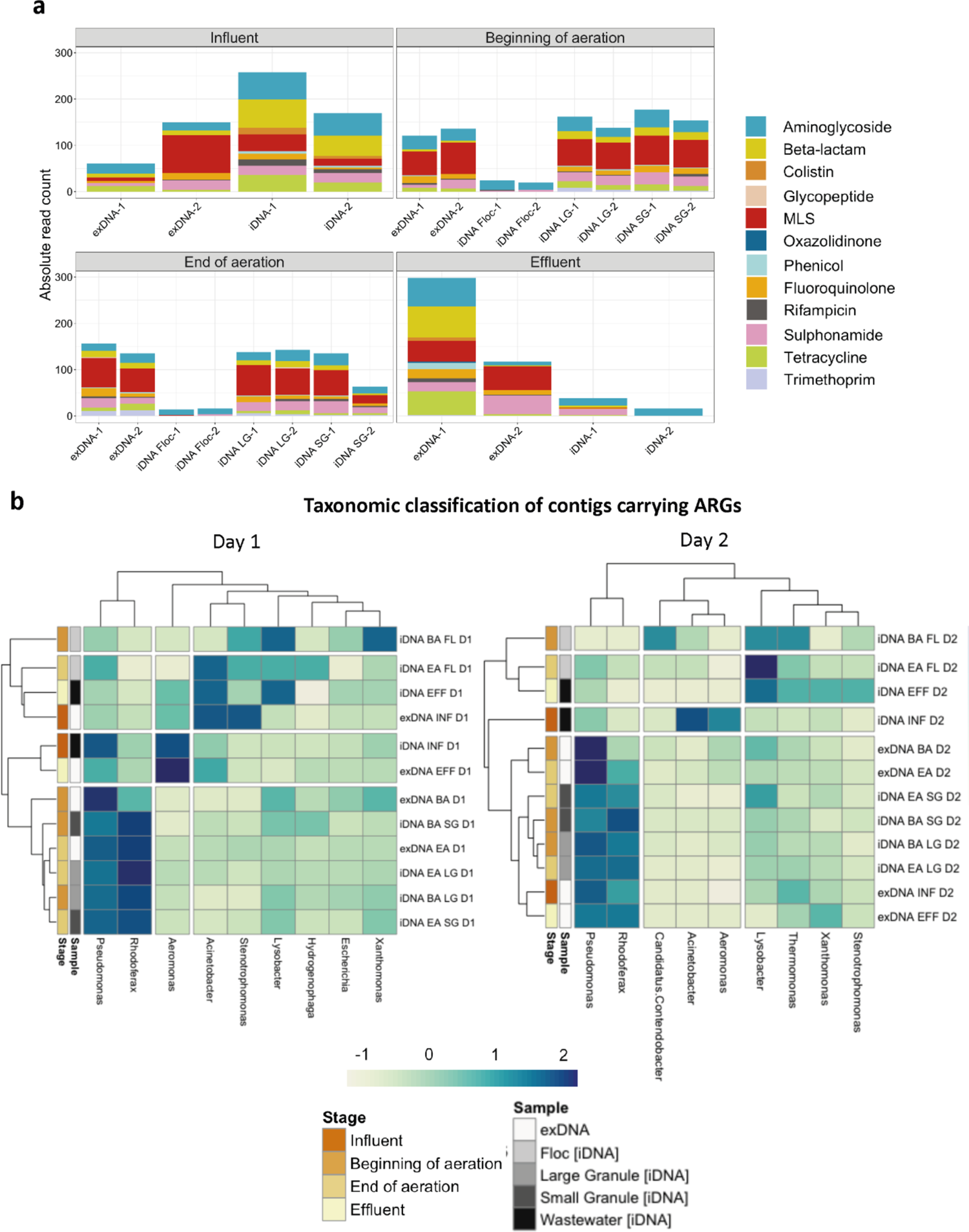
**(a)** Absolute numbers of classified reads of AR-groups. Labels: (F) floccular aggregates, (LG) large granules, (SG) small granules, (−1) first biological replicates, and (−2) second biological replicates. **(b)** Taxonomic classification at genus-level of contigs encoding ARGs heatmap. Colors represent the Z-score computed from the relative abundance at genus-level present above 5% in all classified sequences. Labels: (INF) Influent, (BA) Beginning of the aeration, (EA) End of the aeration, and (EFF) effluent.

All samples, including the different types of bioaggregates, and iDNA and exDNA fractions, contained ARGs. The exDNA resistome profiles at beginning and end of aeration resembled the iDNA resistomes of granules. Similar to observations made on taxonomic affiliations of exDNA sequences, exARGs mainly originated from the granules rather than from flocs. In the effluent, exDNA was a combination of fragments coming from the influent iDNA (**Figure 2b**) (being in the day 2 part of the bigger dendogram) plus the different bioaggregates.

The 5 most abundant ARGs over all samples were: aminoglycoside *arr-3 gene* (n = 58 reads), tetracycline *tet(C)* (n = 46 reads), sulfonamide *sul1* (n = 44 reads), multidrug efflux pump *oqxB* (n = 34 reads), and aminoglycoside *aph(3’’)-Ib* (n = 34 reads). Taken together (n = 216 reads), these genes only amount to 7.6% of the total ARGs identified. This high diversity of resistance genes matches with another metagenomic analysis of activated sludge samples (Zhou et al., 2019).

Of all 478 ARGs identified in the effluent, 89% were carried by exDNA. This can present a risk for AMR dissemination in the environment and for health by natural transformation of exDNA fragments in microorganisms present in receiving water bodies and drinking water resources, especially for the resistance determinants to the last-resource antibiotics such as carbapenem and colistin.

The types of exARG and iARG were similar, matching with Zhou et al. (2019). In contrast, using PCR only, Li et al. (2020) detected dissimilar compositions of iARGs and exARGs during sludge granulation at pilot scale. Early-stage granulation is a dynamic process involving changing community compositions along the establishment of physicochemical gradients (*e.g.*, increasing anaerobic zones in granules) (Sengar et al., 2018). This can lead to bacteria decay and DNA release shifts over the process, therefore leading to variations in ARG profiles as well. Here, mature granules from the full-scale AGS WWTP were used to track the fate of ARGs under pseudo steady-state conditions.

### 3.4. Gram-negative bacteria are potential carriers of ARGs in the AGS

All contigs >500 bp containing ARGs were taxonomically classified to identify which microorganisms carried and potentially released ARGs (**Figure 2b; Figure S4**). As high as 98% of contigs containing ARGs were annotated with taxonomy.

A contig affiliated to a taxon does not necessarily mean that the read comes from that specific organism: different populations can share similar ARG sequences. The contigs containing ARGs over the whole metagenomic dataset of all samples and fractions were affiliated to Gram-negative genera like *Pseudomonas* (8.4 ± 0.7%), *Rhodoferax* (5.6 ± 0.9%), *Acinetobacter* (4.3 ± 1.4%), *Aeromonas* (3.1 ± 1.1%), *Xanthomonas* (3.9 ± 0.9%), and *Acidovorax* (1.9 ± 0.4%) as well as the Gram-positive and antibiotic-producing genus *Streptomyces* (1.6 ± 0.4%) (**Figure S4**).

*Pseudomonas* and *Rhodoferax* genera’s high abundance pattern were similar between the two sampling days (**Figure 2b**). iDNA contigs containing ARGs in small and large granules were affiliated at several instances to *Rhodoferax* (10.1 ± 0.4% in granules). Out of the contigs where ARGs were localized, the ARG-containing exDNA affiliated to *Rhodoferax* (beginning of aeration: 2.0 ± 0.5%; end of aeration: 11.0 ± 2.9%). This increase suggests that DNA was released from *Rhodoferax* cells during aeration. *Rhodoferax* is abundant in WWTP effluents and often identified as AMR carriers by encoding multiple efflux pumps (Jin et al., 2020; Zhou et al., 2020). ARGs have been identified from metagenome-assembled genomes (MAGs) retrieved from activated sludge from Danish WWTPs (Miłobedzka et al., 2022; Singleton et al., 2021): *Rhodoferax* was the most abundant population containing multiple ARGs.

The effluent iDNA is mainly be composed of microorganisms that go through the process without being removed, from bacteria growing on suspended solids and of microorganisms detached from the AGS bioaggregates. Some microorganisms that entered the AGS plant in high abundance and passed through the WWTP are *Aeromonas* and *Acinetobacter,* especially in day 1. The effluent was enriched by microorganisms coming mainly from floccular sludge such as *Lysobacter* (both days) and *Thermomonas* (day 2) (**Figure 2b**). Pathogenic *Arcobacter* crossed WWTPs from influent to effluent (**Figure 1c**) but interestingly, no ARG was detected in any of *Arcobacter* contigs. Further research needs to be done to clearly identify which ARGs in which MGEs are inside specific microbial hosts. Recent analytical advances now allow to identify potential hosts of resistances using Hi-C sequencing (Stalder et al., 2019): populations of *Aeromonadaceae, Moraxellaceae,* and *Bacteroidetes* were shown to be critical reservoirs of AMR in WWTPs.

Collectively, the resistome results highlight that AMR determinants released in exDNA during the WWT operation are importantly linked to Gram-negative bacteria, notably *Rhodoferax,* and to pathogenic bacterial genera carrying ARGs in the effluent such as *Pseudomonas* and *Aeromonas*.

### 3.5. MGEs and ARGs often co-localize on contigs recovered from the exDNA of the effluent

Both iDNA and exDNA fractions and all bioaggregates were sources of a diversity of MGEs. The different samples yielded 55,344 different MGEs belonging to bacterial insertion sequences (9,845 reads), integrons, integrases and gene cassettes (20,149 reads), and bacterial integrative conjugative elements (25,350 reads).

When ARGs and MGEs co-localize on the same genetic fragment, there is an increased chance that the fragment can be transferred between bacterial cells. Since MGEs facilitate the transfer, integration, and transposition of genes in genomes, this poses a clear risk for ARG dissemination. From all contigs >500 bp, 312 contigs were identified to contain both ARGs and MGEs (**Table S6**): their 5 most abundant ARGs were *blaVIM-48* (28), *cmx-1* (28), *sul2* (25), *tetA* (24), and *aadA6* (24). A set of 80 contigs contained multiple ARGs, and of these, 60 also contained MGEs: their 5 most abundant ARGs were all aminoglycoside resistance genes, namely *aph(3’’)-Ib* (17), *aadA6* (17), *aph(6)-Id* (17), *aadA11* (7), and *aac(3)-I*b (7).

Multiple co-localization patterns were detected only in the iDNA of granules and in the exDNA of the effluent. For instance, *sul2* and *tetA* were exclusively found in the iDNA from large granules (replicate 2) from the end of the aeration, and in the effluent exDNA fraction (replicate 2). This contig also aligned with ISVsa3, an insertion sequence often found on plasmids (Toleman and Walsh, 2010).

Of the 60 contigs >500bp were multiple ARGs and MGEs co-localized, 3 major groups of contigs are highlighted hereafter because of their abundance in the system or the risk associated with leaving the WWTP in the exDNA of the effluent.

Firstly, as shown in **Table S6**, 15 contigs were identified both as carriers of two aminoglycoside resistance genes (*aph(3’’)-Ib* and *aph(6)-Id)*, and as part of transposon Tn5393 (**Figure 3a**). These contigs were found in all samples across WWT operation for all microbial aggregates and exDNA and iDNA fractions (excluding the flocs). After alignment to the NCBI nucleotide database, the contigs aligned with IncQ plasmids, mostly from populations of *Acinetobacter* and *Pseudomonas* genera, previously identified in our wastewater samples. IncQ plasmids are significant since they are related to the broadest host range of all replicating elements in bacteria (Meyer, 2009). They are known to mobilize via conjugation although natural transformation has also been observed (Bönemann et al., 2006; Meyer, 2009; Szczepanowski et al., 2009, 2004).

**Figure 3.**
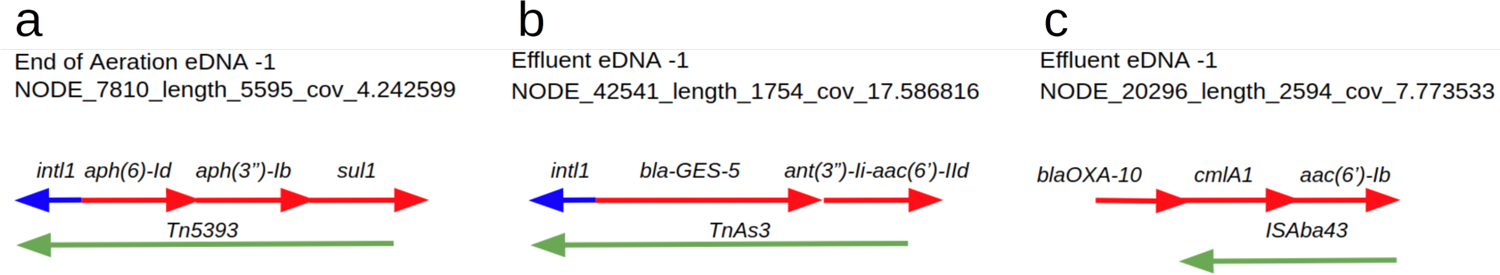
Contigs containing both ARGs and MGEs. **(a)** Contig of length 5595 bp, containing the class 1 integron-integrase (*intI1*), aminoglycoside phosphotransferase (*aph(6-Id),* also known as *strB*), an aminoglycoside phosphotransferase (*aph(3’’)-Ib* also known as *strA*), a sulfonamide resistant dihydropteroate synthase (*sul1*), and a fully classified transposon Tn5393. **(b)** Contig of length 1754 bp, containing the class 1 integron-integrase (*intI1*), beta-lactamase *bla-GES-5*, integron-encoded aminoglycoside acetyltransferase fusion protein *ant(3’’)-li-aac(6’’)-IId*, fully classifying as transposon TnAs3. **(c)** Contig of length 2594 bp, containing beta-lactamase *blaOXA-10*, a plasmid or transposon-encoded chloramphenicol exporter gene *cmlA1*, an integron-encoded aminoglycoside acetyltransferase *aac(6’)-Ib,* and a transposase ISAba43.

Then, 18 contigs aligned with a Tn3 family transposon (TnAs3), normally encoding a β-lactamase gene, a transposase and a resolvase (Nicolas et al., 2015) as identified from the exDNA effluent fraction (**Figure 3b**). These contigs (identities >95%) aligned exclusively with plasmids from *Escherichia coli, Klebsiella pneumoniae, Pseudomonas aeruginosa, Enterobacter cloacae, Enterobacter kobei,* and *Klebsiella oxytoca*. Interestingly, of all pathogens, the World Health Organization stressed that addressing carbapenem-resistance in *Pseudomonas aeruginosa* and *Enterobacteriaceae* should have the highest priority (Tacconelli and Magrini, 2017).

Finally, another contig of importance found in the exDNA fraction of the effluent carried multiple resistance genes against aminoglycosides (*aac(6’))*, beta-lactamases (*blaOXA-10)*, and phenicols (*cmlA1)* (**Figure 3c**). This contig aligned with a 100% sequence identity to a plasmid already annotated from *Aeromonas hydrophila* (pWCX23_1), carrying 15 different ARGs (Nwaiwu and Aduba, 2020).

The strong presence of mobilized resistance determinants in the extracellular fraction of the AGS WWTP has been also described by Ikuma and Rehmann (2020). Using mathematical modelling, they suggested that, at the WWTP effluent discharge point in a river, the total number count of ARGs was 13 times higher when including exARGs on top of iARGs. Collectively, exARGs in WWTP discharges need to be taken into account for accurate risk assessments of AMR.

Co-localization studies are valuable tools to identify the genetic structures and potential hosts by which resistance determinants are carried and transferred between bacteria. More research needs to be conducted to reconstruct full plasmids from wastewater effluents. This will shed light on the type of plasmids released into the environment, *i.e.*, narrow/broad-range and synthetic/naturally occurring, and on the genetic constructions ARGs are embedded in.

### 3.6. No antibiotic-related biosynthetic gene clusters were found in the biomass for the most abundant ARG identified

Functional and pathway analyses were performed on the contigs to identify AMR pathways and on the MAGs to identify potential intrinsic antibiotic production by microorganisms that could promote the development and survival of ARB in AGS systems.

From all the AMR pathways available in the KEGG database (Kanehisa et al., 2002), we identified the cationic antimicrobial peptide (CAMP) resistance pathway, vancomycin resistance pathway, and beta-lactamase resistance pathway, including efflux pumps that actively transport the beta-lactam antibiotics out of the cell (RND efflux pumps) (**Figure S6-S8**). Both influent and effluent iDNA and exDNA were sources of genes involved in beta-lactam resistance. Most genes involved in these resistance pathways were present in the influent iDNA but not in the influent exDNA. However, in the effluent, both iDNA and exDNA contained resistance pathways genes.

From the antiSMASH analysis of MAGs recovered from small and large granules (no MAGs could be recovered from flocs due to the sequencing depth and short reads used) at the end of aeration, positive hits on non-ribosomal peptide synthetase (NRPS) biosynthetic gene clusters (BGCs) were detected in MAGs of *Nitrospira* and *Janthinobacterium*. NRPS antibiotics include vancomycin, polymyxin, and teixobactin, which are highly effective against multidrug-resistant bacteria (Li et al., 2018). The vancomycin resistance pathway was one of the few reconstructed pathways in the samples analyzed (**Figure S6**). Terpenes, indoles, T1PKS, and ectoine BGCs were identified as the most recurrent (**Table S8**). No aminoglycosides or beta-lactams antibiotic BGCs were found in any of the DNA samples.

### 3.7. ARGs increase in exDNA over the WWT operation

To quantify the ARG and ARB occurrence and related removal capacity of the AGS WWTP, a panel of six ARGs and *intI1* as MGE were quantified by qPCR. Two normalization methods (using *16S rRNA* and *rpoB* genes) were assessed. Both the ARGs and the MGE were consistently detected across all the AGS process phases and bioaggregates.

The most predominant genes throughout the process were the sulfonamide resistance gene *sul1* and the class 1 integron gene *intI1* (**Figure 4a**). These genes were the most concentrated ones in the effluent with 7.5 (*sul1*) and 7.0 (*intI1*) log_10_ gene copies mL^-1^ (sum of iDNA and exDNA pools). Genes with the lowest concentration were *bla_CTXM_* (2.1), *qnrS* (3.1), and *tetO* (3.2) (same units as above). Absolute and normalized concentrations of ARGs and MGEs, and iDNA *vs*. exDNA analysis, are shown in **Figure S9** and **Table S7**. These ARGs concentrations are in agreement with recent studies (Calderón-Franco et al., 2020; Pallares-Vega et al., 2019) and comparable to others, especially when considering the variance in sewage and treatment efficiency (Alexander et al., 2020; Lee et al., 2017; Wang et al., 2020).

**Figure 4.**
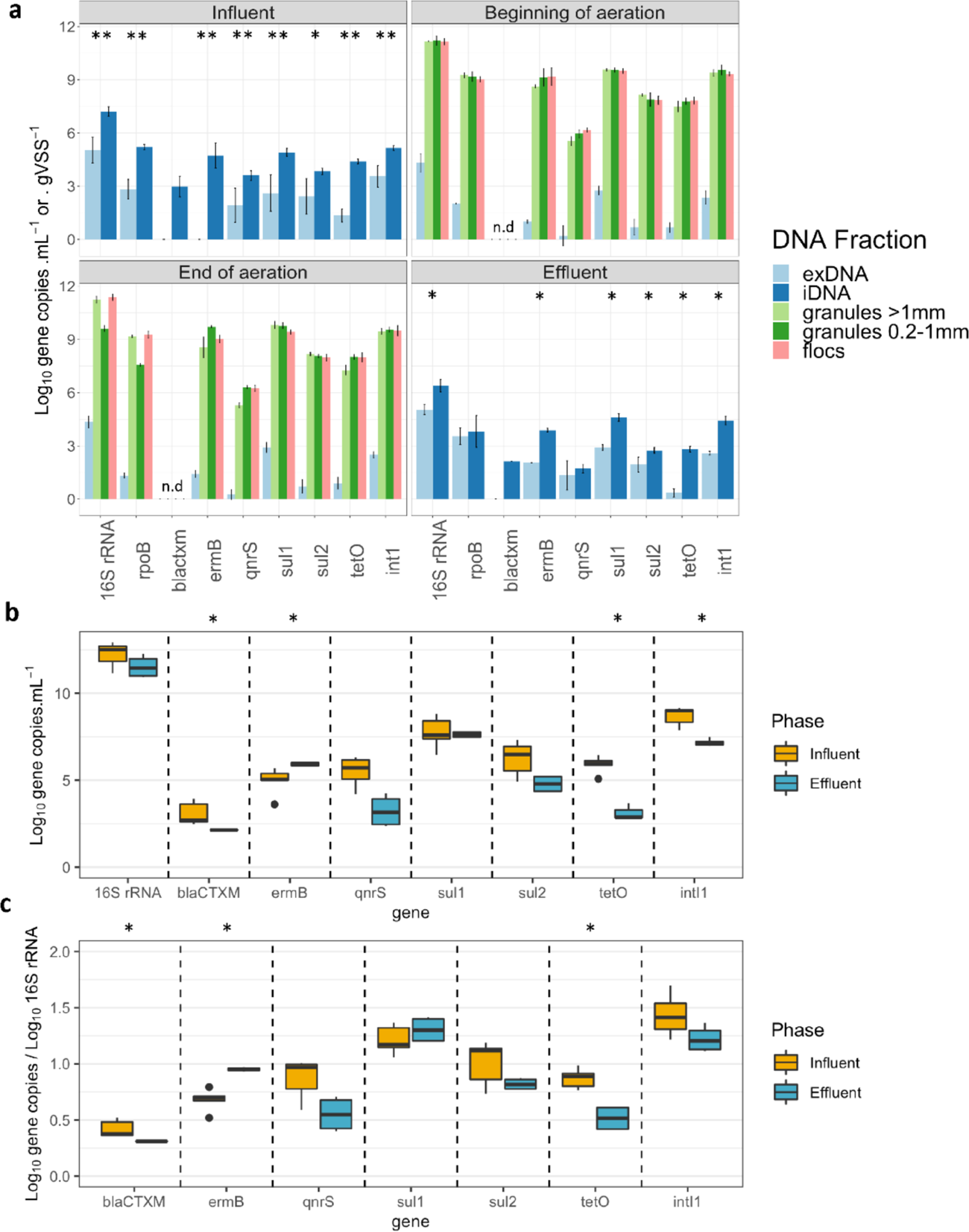
**(a)** Concentration of selected ARGs and *int1I* across AGS WWT phases. Error bars represent the standard deviation between biological replicates (n=3). The significances of the difference between exDNA and iDNA measurements are indicated by the Wilcoxon test and displayed as p<0.05 (*) and p<0.005 (**). **(b)** Absolute abundance of ARGs and *intI1* before and after treatment (influent and effluent phase) expressed in log10 gene copies mL^-1^. **(c)** Relative abundance of genes in the influent and effluent expressed as log10 gene copies / log10 16S rRNA copies. The significance of the difference between influent and effluent measurements, as indicated by the Wilcoxon test, is displayed as p<0.05 (*).

The extended-spectrum beta-lactamase (ESBL) gene (bla_CTXM_) was detected only in the influent (3.0 ±

0.6 log_10_ gene copies mL^-1^) and effluent (2.1 ± 0.1 log_10_ gene copies mL^-1^) iDNAs. No copies of this gene were found during the aeration process or in the exDNA fractions. In the most recent NethMap/MARAN report presenting data on antibiotics use and resistance in the Netherlands (RIVM and SWAB, 2020), the increasing detection of bacteria producing ESBL, even in countries with non-extensive use of antibiotics, was highlighted as an imminent threat. The low but increasing presence of ESBL-producing bacteria can justify the small but significant detection of beta-lactam variants such as bla_CTXM_.

### 3.8. AGS treatment is efficient at reducing the load of resistance determinants

The absolute concentration of iARGs was significantly higher (p<0.05-0.005) than of exARGs across all SBR phases and bioaggreagates (**Figure 4a**). The average iDNA/exDNA ratio was 1.90 in the influent and 2.3 in the effluent (**Figure S9**). These differences in the concentration of iARGs and exARGs match with a recent report of highest relative abundance of sulfonamide, tetracycline resistance, and integron genes in the iDNA than in the exDNA of an AGS WWTP (Li et al., 2020). We found a significant reduction (p<0.005) after treatment for all genes analyzed by qPCR in the iDNA fraction. In contrast, multiple genes such as the bacterial proxy genes *16S rRNA* and *rpoB* as well as the resistance genes *ermB* (2.1 log_10_ gene copies mL^-1^ higher) and *sul1* (1.0 log_10_ gene copies mL^-1^ higher) increase from the influent to effluent exDNA (**Figure S11**). Di Cesare et al. (2016) have reported different ARGs patterns of exARGs, iARGs, and integrase MGEs: microbial cell decay and cell lysis can potentially release genetic material extracellularly. The free-floating exDNA is subjected to the fluctuating conditions of the WWTP environment differently than iDNA (Li et al., 2020). Its ability to maintain mobility and be involved in resistance spread needs to be studied in the future.

Small granules accumulated higher concentrations of resistance genes (average 0.82 ± 0.31) log_10_ gene copies/ log_10_ *16S rRNA* copies) in their iDNA than big granules (0.62 ± 0.26) or flocs (0.61 ± 0.25) at the end of the aeration. No significant differences were observed between granules at the beginning of the aeration (**Figure 4a; Figure S9**). It is not clear why small granules accumulated more ARGs during the 9-h aeration process.

Flocs and small granules are more susceptible to immigration from the influent/sewage than large granules (Ali et al., 2019). The solid retention time (SRT) of small granules is much shorter (7.7 ± 0.5 days) than that of big granules (142.6 ± 14.9 days) (Ali et al., 2019). If the influent is enriched with ARB, immigration can explain a faster loading of ARGs in small than large granules. By being more rapidly released out of the tank, small granules can play a role in the release and dissemination of ARGs through WWTP effluent than big granules. Because of their higher specific surface area, fractions of small granules can capture more free-floating exDNA released from cell decay during the treatment process. exDNA can help stabilize their architecture by adhesion at initial granulation stages through acid-base interactions (Wang et al., 2020; Das et al., 2011, 2010).

Further studies on ARG distribution across bioaggregates are needed to determine if ARGs are homogeneously distributed across the cross-section of the granules or mainly located at their surface or in one of their layers. Fluorescent *in situ* hybridization with ARG probes could determine the ARG distribution in granule slices. Such studies will also indicate if transfer between ARGs is more prone to happen between or within granules.

### 3.9. Resistance loads in wastewater effluents depend on DNA fraction

Looking at the removal capacity of the full process from influent to effluent, the overall pool of resistance determinants was significantly reduced during WWT (average 1.1 log_10_ gene copies mL^-1^ removal). Some ARGs were still discharged with iDNA inside cells (2-4 log_10_ gene copies g VSS^-1^) and with exDNA free-floating in water (0.5-3 log_10_ gene copies mL^-1^) (**Figure 4a; Figure S9**). The majority of the targeted genes decreased after the treatment. From the panel analyzed by qPCR, *intI1, bla_CTXM_*, and *tetO* significantly (p<0.05) decreased by 1.3, 1.7, and 2.3 log_10_ gene copies mL^-1^, respectively. The AGS process is partially effective at reducing ARGs: Sabri et al. (2020) have reported a similar drop for tetracycline resistance gene (*tetW*) in an AGS technology of around 2 log_10_ gene copies mL^-1^.

Based on the remaining ARGs concentrations in the effluent, an average of 6.3 log_10_ gene copies mL^-1^ were released in total, iDNA plus exDNA (**Figure 4b**). Taking hospital discharges as references, this can be considered high (>10^4^ gene copies mL^−1^) (Le et al., 2016). From **Figure 4c**, the normalized values did not significantly enrich bacterial populations carrying ARGs or integrases.

Overall, we found that the AGS WWTP can reduce the amount of ARB but still releases in the environment a significant amount of ARGs enclosed on MGEs of free-floating exDNA. The commonly overlooked exDNA should therefore be considered (together with the iDNA of ARB) as an important factor in the dissemination of resistance determinants in the aquatic environment. Risks associated with exDNA and iDNA fractions discharged in water resources need to be evaluated, with a clear identification of exposures and effects.

## 4. Conclusions

This work led to the following main conclusions:

1. *Pseudomonas* and *Rhodoferax* populations were the main hosts of antibiotic resistance genes (ARGs) in the biomass. Other ARG-carrying populations affiliated with *Acinetobacter*, *Aeromonas*, *Xanthomonas*, *Acidovorax, Bacteroidetes,* and *Streptomyces*.
2. Several ARGs co-localized with mobile genetic elements (MGEs) on genetic contigs of free-floating extracellular DNA, thus being potentially transferrable to microorganisms. The most abundant ARGs co-localizing with MGEs were the beta-lactamases resistance gene *blaVIM-48*, chloramphenicol resistance gene *cmx-1*, sulfonamides resistance gene *sul2*, tetracycline resistance gene *tetA*, and aminoglycoside resistance gene *aadA6*.
3. The panel of ARGs and *intI1* MGE analyzed by qPCR were detected in all samples.
4. The ARG fraction in intracellular DNA decreased with 1.1 log_10_ gene copies mL^-1^ during the process. Conversely, some ARGs located in the exDNA (*ermB, sul1* and *sul2*) increased during the process.
5. According to metagenomics, exDNA carries ARGs enclosed inside a diversity of MGEs, detected in the WWTP effluent. These can potentially spread AMR in aquatic environments by horizontal gene transfer.

Therefore, studies for the surveillance, risk assessment, and mitigation of AMR in wastewater environments should consider not only iDNA but also exDNA pools.

## Data availability

Metagenome sequencing data were deposited in the NCBI database with the BioProject ID PRJNA783874.

## Authors’ contributions

DCF designed the study with inputs of MvL, MP, TA, and DGW. DCF, RS, and SC performed the experimental investigations. DCF wrote the manuscript with direct contribution, edits, and critical feedback by all authors.

## Acknowledgements

We are grateful to Pascalle Vermeulen from Royal HaskoningDHV, The Netherlands, for helping us arrange and design this sampling campaign. This work is part of the research project “Transmission of Antimicrobial Resistance Genes and Engineered DNA from Transgenic Biosystems in Nature” (TARGETBIO) funded by the programme Biotechnology & Safety Program of the Ministry of Infrastructure and Water Management (grant no. 15812) of the Applied and Engineering Sciences (TTW) Division of the Dutch Research Council (NWO).

## List of abbreviations

aac(3): Ib Aminoglycoside 3’-N-acetyltransferase resistance gene

AMR: Antimicrobial resistance

*aadA6*: Aminoglycoside (3’’) adenyltransferase resistance gene

*aadA11*: Aminoglycoside (3’’) (9) adenyltransferase resistance gene

AGS: Aerobic granular sludge

*aph(3”)-Ib*: Aminoglycoside 3’-phosphotransferase resistance gene

*aph(6)-Id*: Aminoglycoside O-phosphotransferase resistance gene

ARB: Antibiotic-resistant bacteria

ARG: Antibiotic resistance gene

*arr-3*: Plasmid-encoded ribosyltransferase resistance gene

BGC: Biosynthetic gene clusters

COG: Clusters of Orthologous Genes

exARG: Free-floating extracellular antibiotic resistance gene

exDNA: Free-floating extracellular DNA

HGT: Horizontal gene transfer

iARG: Intracellular antibiotic resistance gene

iDNA: Intracellular DNA

MAG: Metagenome-assembled genome

MGE: Mobile genetic element

*oqxB*: Fluoroquinolone efflux pump membrane transporter gene

qPCR: Quantitative polymerase chain reaction

SBR: Sequential batch reactor

*sul1*: Sulfonamide resistant dihydropteroate synthase of Gram-bacteria, linked to class 1 integrons

*tetC*: Tetracycline efflux pump gene

WWT: Wastewater treatment

WWTP: Wastewater treatment plant

## Supplementary material

### Quantitative PCR conditions, primers and standards

Quantitative polymerase chain reaction experiments were performed for the quantification of the genes in **Table S1** in the WWTP’s DNA samples (Section 2.1) using the thermal cycler qTOWER3 (Analytica Jena, Germany). Based on the measured cycle threshold values (Ct values), standard curves were derived, which indicated the efficiency of the reaction and enabled further determination of the genes’ concentration. The DNA standards sequences, the standard curves, the reaction results per qPCR, and the calculation procedure are mentioned in Appendix A. The primers used for each gene and the amplified fragments’ lengths are listed in **Table S1**. The qPCR analysis was performed in a total volume of 20 μL using 10 μL iQ™ SYBR® Green Supermix1x (BIO-RAD Laboratories; containing dNTPs, iTaq™ DNA Polymerase, MgCl_2_, SYBR® Green I, enhancers, stabilizers, and passive reference dye fluorescein), 0.2 μL of each primer (10 µM), 2 μL DNA, and 7.6 μL Molecular Biology grade water (Sigma Aldrich, USA). Three technical replicates of each sample were used in all cases to improve accuracy and precision. The thermal cycler’s amplification conditions were adjusted according to each gene’s properties, and different annealing conditions were applied based on primers’ melting temperatures.

**Table S1.**
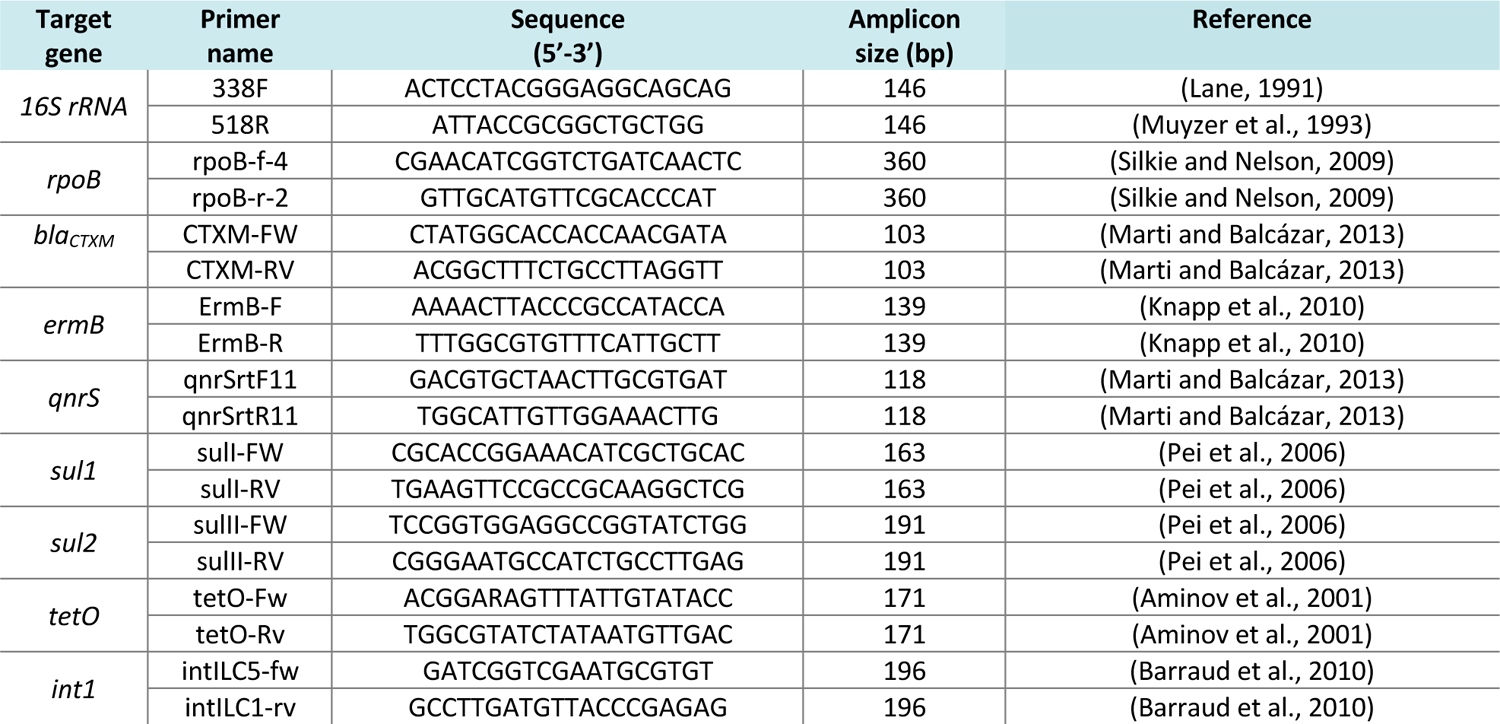
Primer sequences used for qPCR amplification in the study and their respective amplification sizes.

**Table S2.** **qPCR thermal cycler conditions.** The temperature and duration of the annealing phase were different for each gene: ***qnrS, ermB, sul1, bla_CTXM_*** – 60°C, 30 sec; ***int1, tetO*** – 60°C, 1 min; ***sul2*** – 61°C, 1 min; ***16S rRNA*** – 55°C, 20 sec; ***rpoB*** – 55°C, 30 sec.

| Genes | Step | T (°C) | Time | Genes | Step | T (°C) | Time |
| --- | --- | --- | --- | --- | --- | --- | --- |
| *qnrS*, *sul2*,<br>*ermB*,<br>*bla*<sub>CTXM</sub> | 1 | 95 | 5 min | *16S rRNA*,<br>*rpoB*, *tetO*,<br>*sul1*, *int1* | 1 | 95 | 10 min |
|  | 2 | 95 | 15 sec |  | 2 | 95 | 15 sec |
|  | 3 | Mentioned in description |  |  | 3 | Mentioned in description |  |
|  | 4 | 72 | 30 sec |  | 4 | 72 | 10 sec |
|  | 5 | 80 | 5 sec |  | 5 | 80 | 1 sec |
|  | 6 | 80 | 1 sec |  | 6 | 95 | 15 sec |
|  | 7 | 12 | 59 sec |  | 7 | 60 | 1 min |
|  | 8 | 12 | 1 sec |  | 8 | 95 | 15 sec |

**Table S3.**
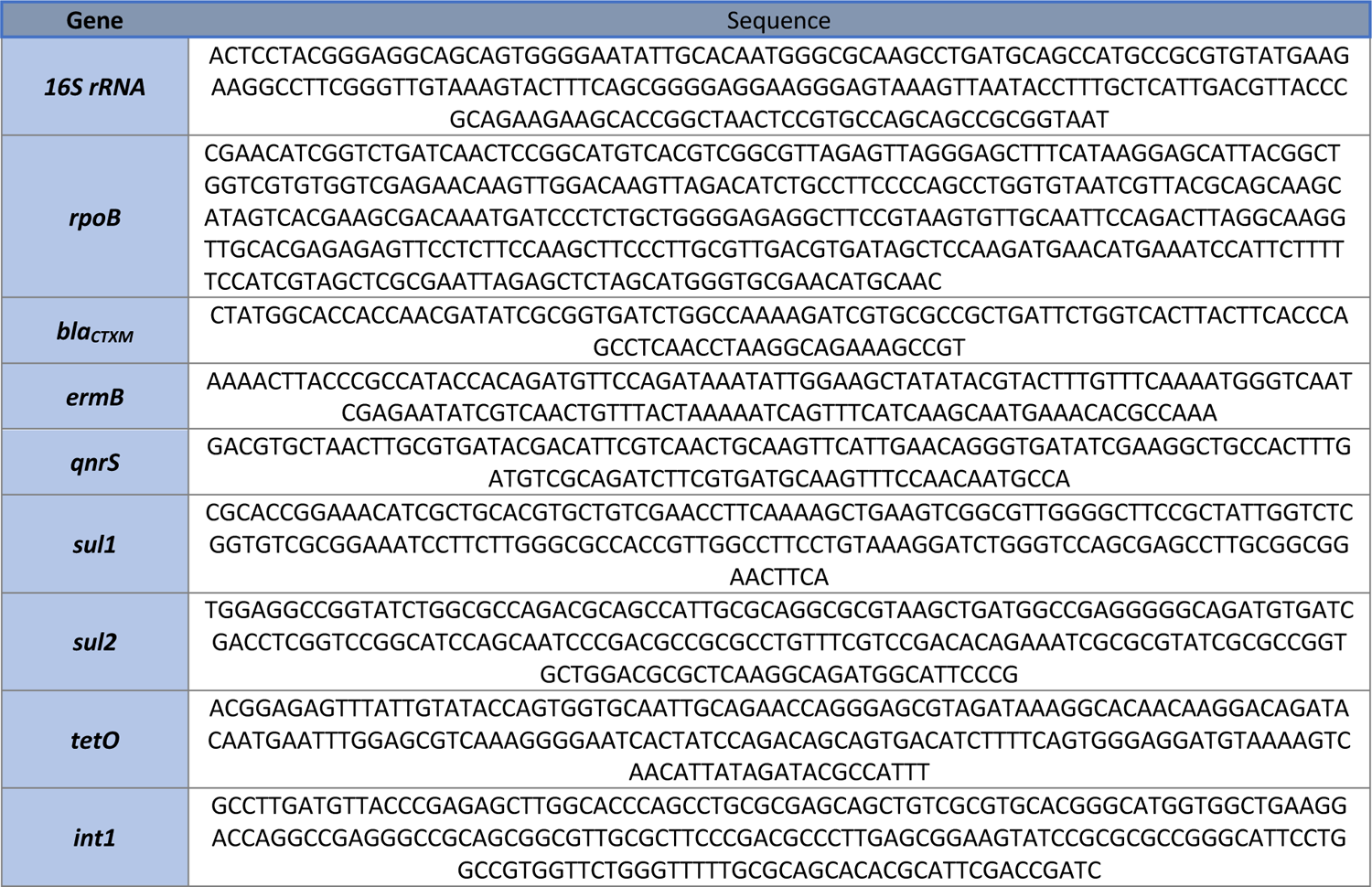
Synthetic DNA fragments (gBlocks) used for generating standard curves.

### Quantitative PCR statistical results

**Table S4.** P-values indicating the statistical significance of the difference between the genes’ absolute concentration in the intracellular and the extracellular DNA from influent and the effluent samples.

| Gene | Influent | Effluent |
| --- | --- | --- |
| *16S rRNA* | 0.005 | 0.0294 |
| *bla<sub>CTXM</sub>* | n/a | 0.1939 |
| *ermB* | 0.003 | 0.02107 |
| *qnrS* | 0.005 | 0.3768 |
| *sul1* | 0.005 | 0.03038 |
| *sul2* | 0.013 | 0.03038 |
| *tetO* | 0.005 | 0.0294 |
| *int1* | 0.005 | 0.0294 |
| *rpoB* | 0.005 | 0.376 |

**Table S5.** P-values indicating the statistical significance of the differences between the genes’ absolute concentration in the different microbial aggregates that constitute the AGS process. LG = Large granules > 1mm, SG = Small granules < 1 mm.

| BEGINNING OF AERATION |  |  |  |
| --- | --- | --- | --- |
| Gene | LG vs SG | BG vs flocs | SG vs flocs |
| *16S rRNA* | 0.030 | 0.061 | 0.810 |
| *bla<sub>CTXM</sub>* | n/a | n/a | n/a |
| *ermB* | 1.000 | 1.000 | 0.661 |
| *qnrS* | 0.312 | 0.030 | 0.194 |
| *sul1* | 0.471 | 0.885 | 0.885 |
| *sul2* | 0.665 | 0.030 | 1.000 |
| *tetO* | 0.312 | 0.312 | 0.665 |
| *int1* | 0.471 | 0.885 | 0.665 |
| *rpoB* | 0.381 | 0.312 | 0.059 |
| END OF AERATION |  |  |  |
| Gene | LG vs SG | BG vs flocs | SG vs flocs |
| *16S rRNA* | 0.005 | 0.379 | 0.005 |
| *bla<sub>CTXM</sub>* | n/a | n/a | n/a |
| *ermB* | 0.028 | 0.468 | 0.029 |
| *qnrS* | 0.005 | 0.005 | 0.575 |
| *sul1* | 0.936 | 0.005 | 0.008 |
| *sul2* | 0.066 | 0.045 | 0.936 |
| *tetO* | 0.005 | 0.013 | 0.810 |
| *int1* | 0.297 | 0.936 | 0.575 |
| *rpoB* | 0.005 | 0.574 | 0.005 |

### Metagenomics procedure and quality control

**Figure S1.**
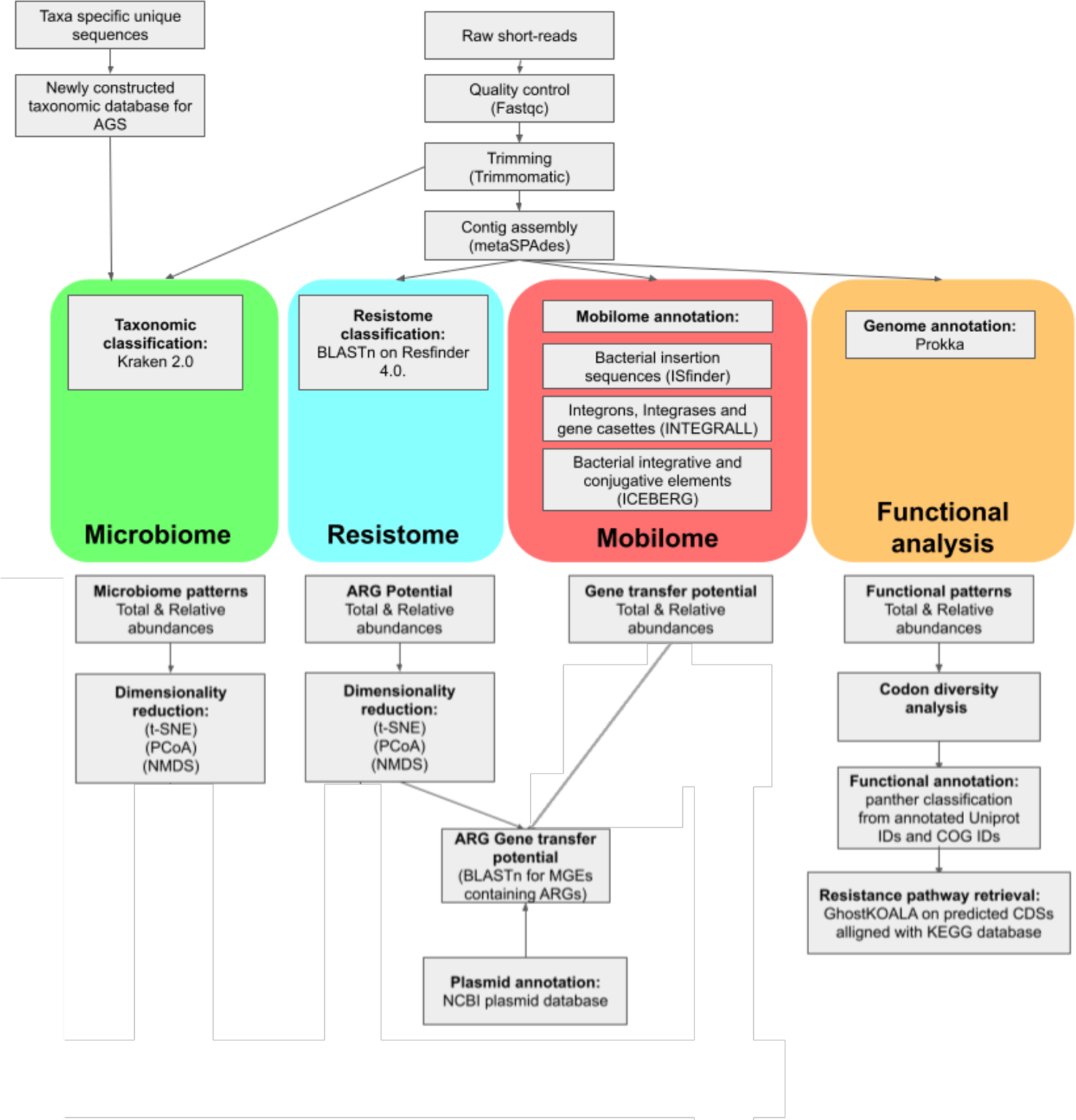
Schematic representation of the procedures followed for the analysis of the metagenomics dataset.

**Figure S2.**
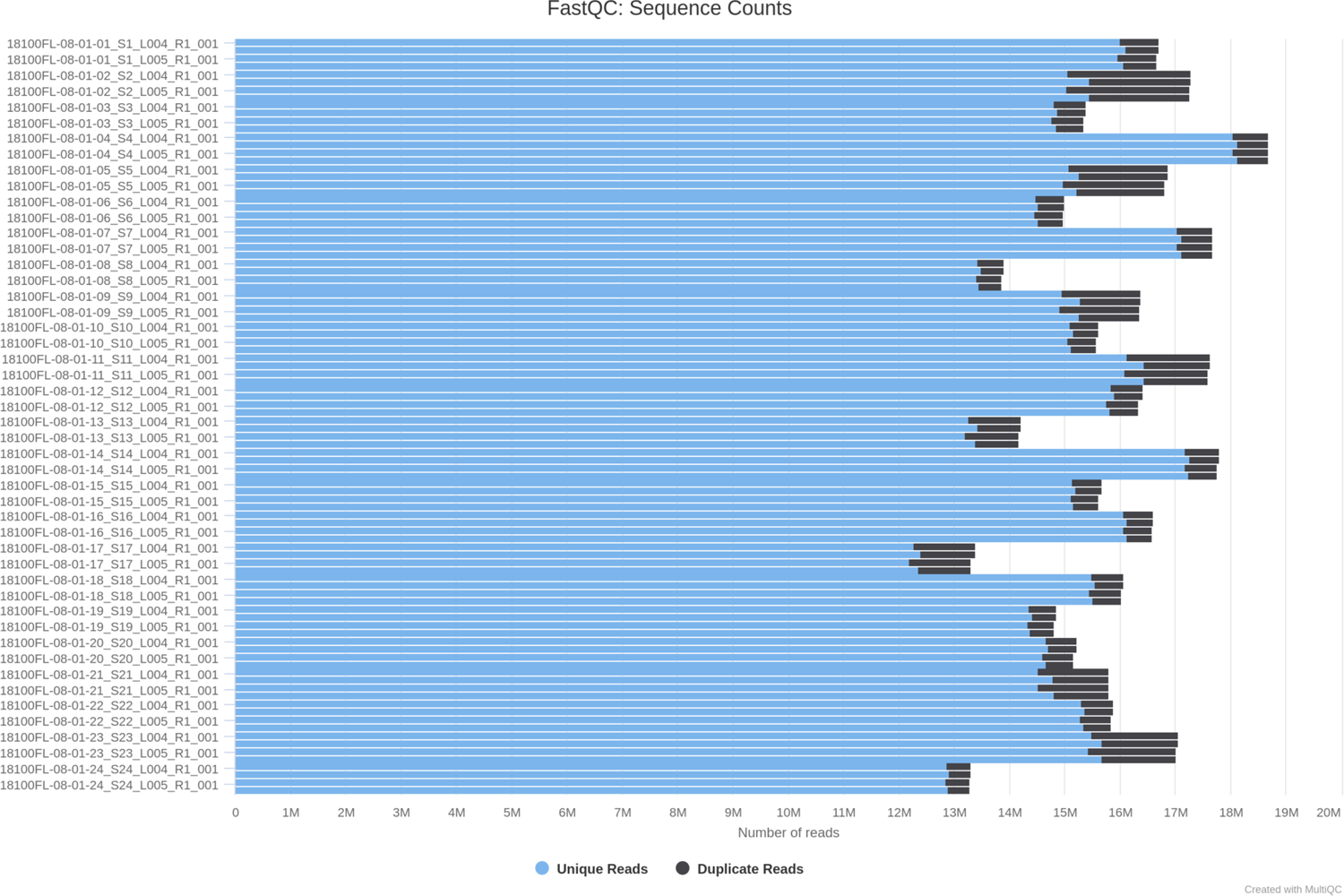
Cumulative sequence counts of the raw Illumina paired-end reads showing the unique reads in blue and the duplicated reads in black.

### Dimension reduction of metagenomics datasets

**Figure S3.**
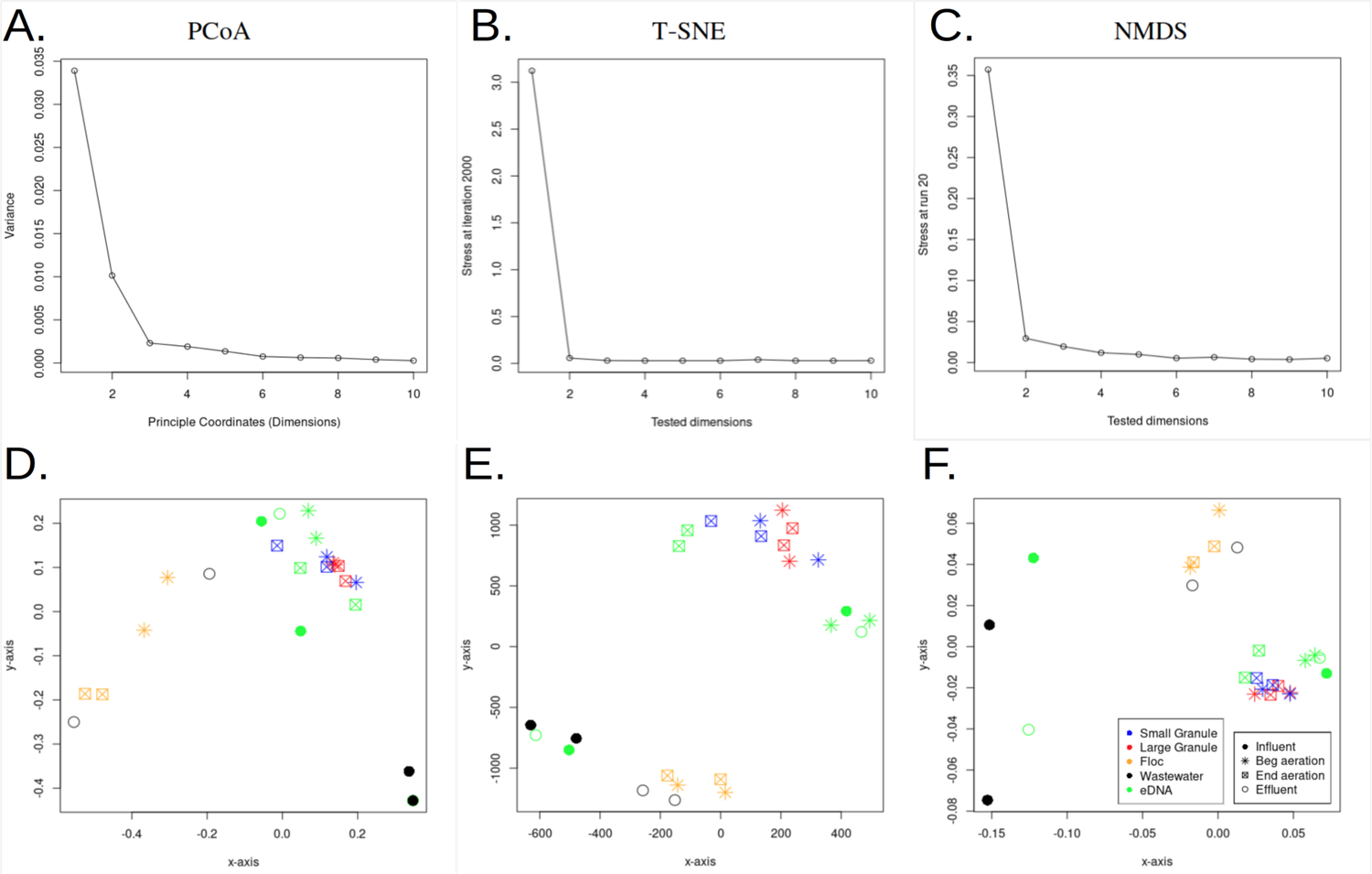
Dimension reduction of the absolute microbiome composition on genus-level after normalization and variance stabilization transformation, with Bray-Curtis as distance metric in all methods. (**A)** shows a scree plot based on PCoA, showing the proportion of variance in every principal coordinate. **(B)** and **(C)** show the stress at the final iteration over ten tested dimensions for t-SNE and NMDS, respectively. **(D)**, **(E)** and **(F)** show the dimension reductions for PCoA, t-SNE, and NMDS, respectively. Colours indicate types of samples: small granule (blue), large granule (red), floc (orange), wastewater (black), and exDNA (green). Shapes indicate stages in the WWT operation: influent (filled circle), beginning of aeration (asterisk), end of aeration (crossed square) and effluent (empty circle).

### Taxonomic classification of contigs from all AGS stages and DNA fractions

**Figure S4.**
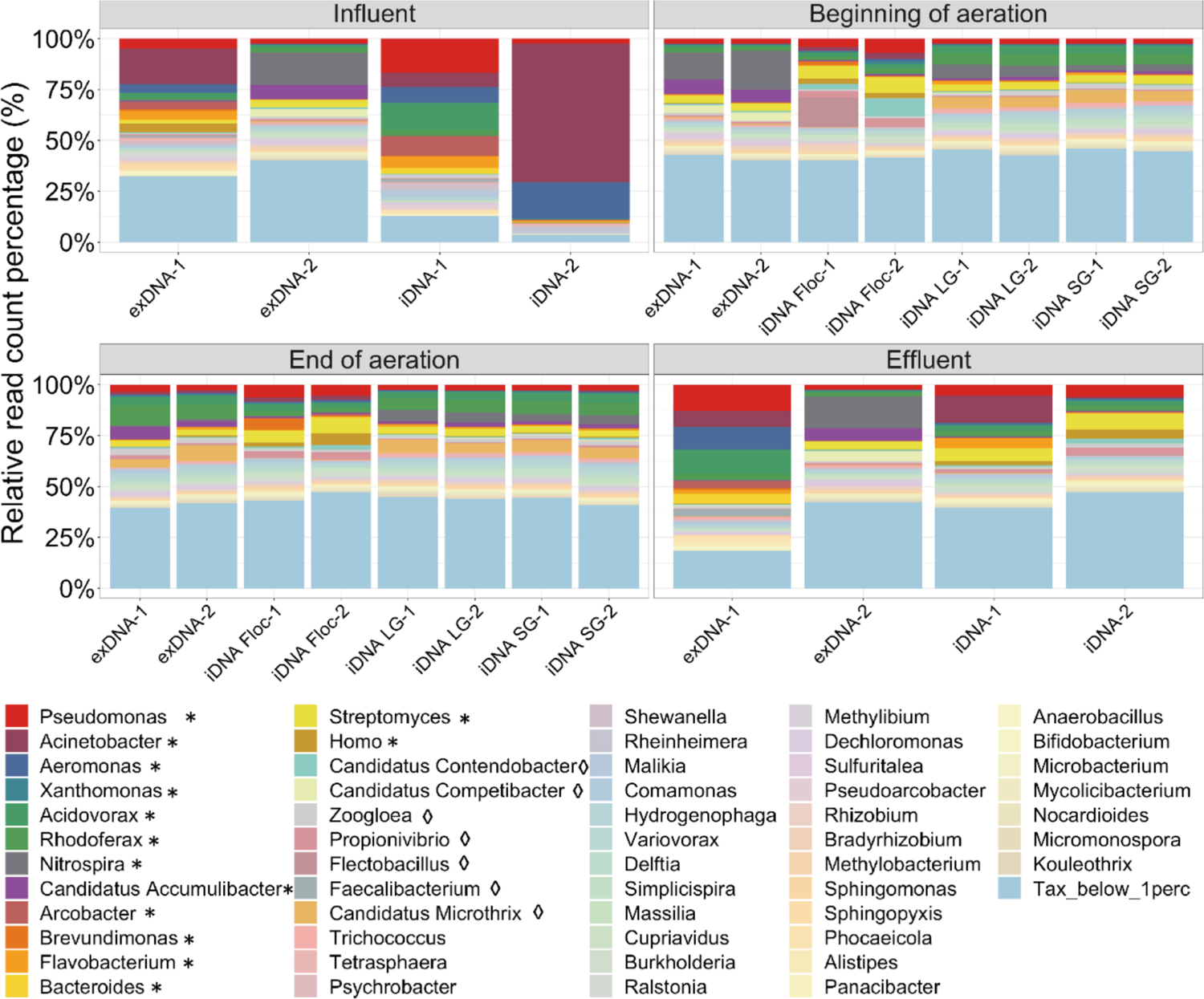
Taxonomic classification at genus-level of short-reads for different DNA fractions and different microbial aggregate types across four stages of AGS WWT operation. A newly adapted database was used to determine relative abundance (%) of classified reads on genus-level within a sample. Labels: (F) floccular aggregates, (LG) large granules, (SG) small granules, (−1) first biological replicates, and (−2) second biological replicates. Asterisk (*) after genera names indicates genera with >3% relative abundance in at least one of the samples, also with the standard Kraken 2.0 database. Diamond (◊) after genera names indicates genera with >3% relative abundance in at least one of the samples, but not with the standard Kraken 2.0 database. Genera with <1% relative abundance in all samples are labelled as “Tax_below_1perc” (light-blue).

### Taxonomic classification of ARGs contigs from all AGS stages and DNA fractions

**Figure S5.**
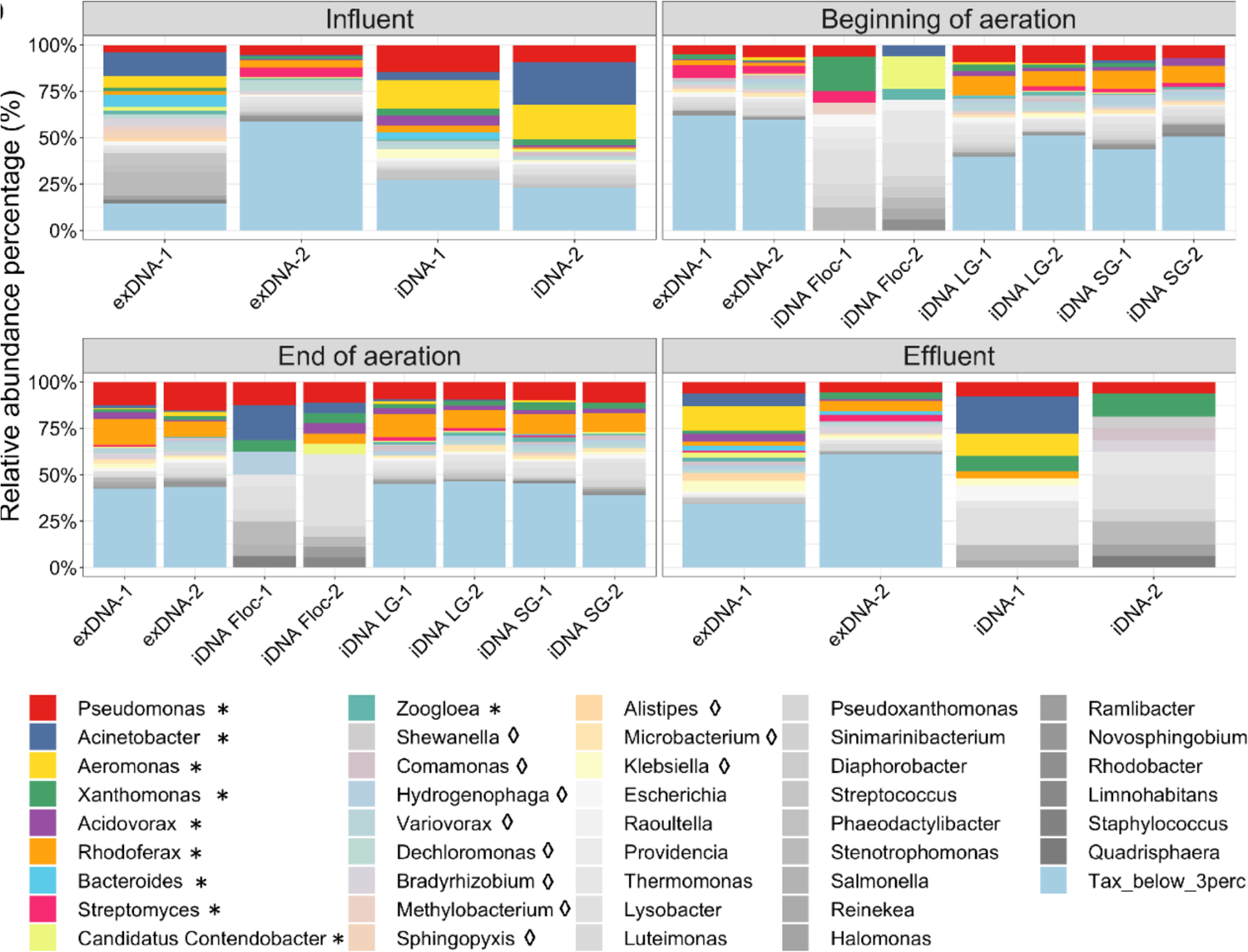
Relative abundance of contigs containing ARGs across the AGS WWT operation annotated at genus-level using the newly constructed database. Bright colors (*) represent genera above 3% that were already present above 3% relative abundance in the general taxonomic classification (**Figure S3**). Pastel colors (◊) represent genera above 3% that were identified below 3% in the general taxonomic classification. Grey-scale colors represent genera above 3% but identified below 1% in Figure 1c. Genera with <3% relative abundance in all samples are labelled as “Tax_below_3perc” (light-blue).

### Reconstructed antibiotic resistance pathways from the metagenomic datasets

**Figure S6.**
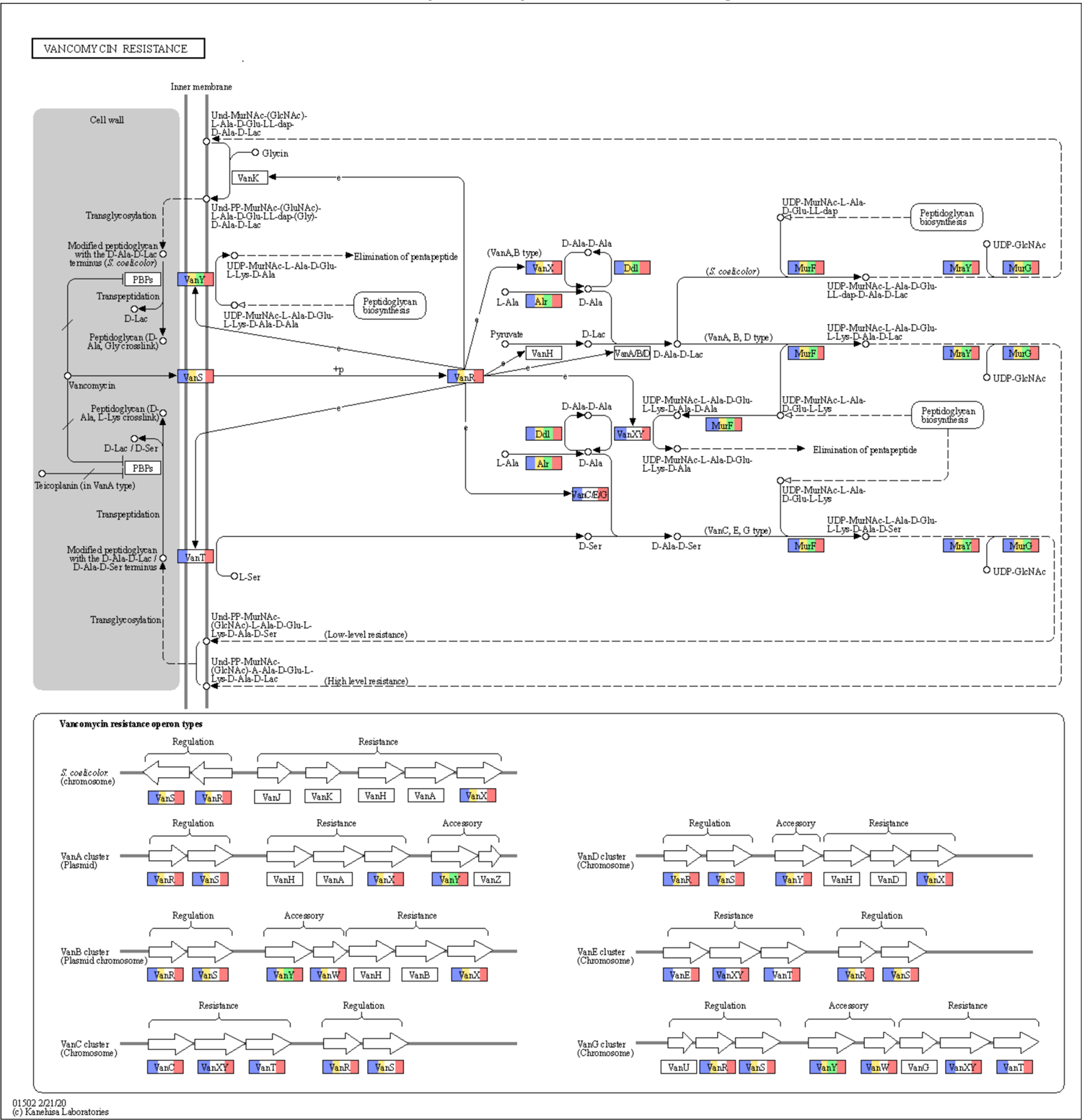
Vancomycin resistance pathway based on KEGG orthology. Colors indicate the genes that were found to be present in a certain sample: influent iDNA (blue), influent exDNA (yellow), effluent iDNA (green), and effluent exDNA (red).

**Figure S7.**
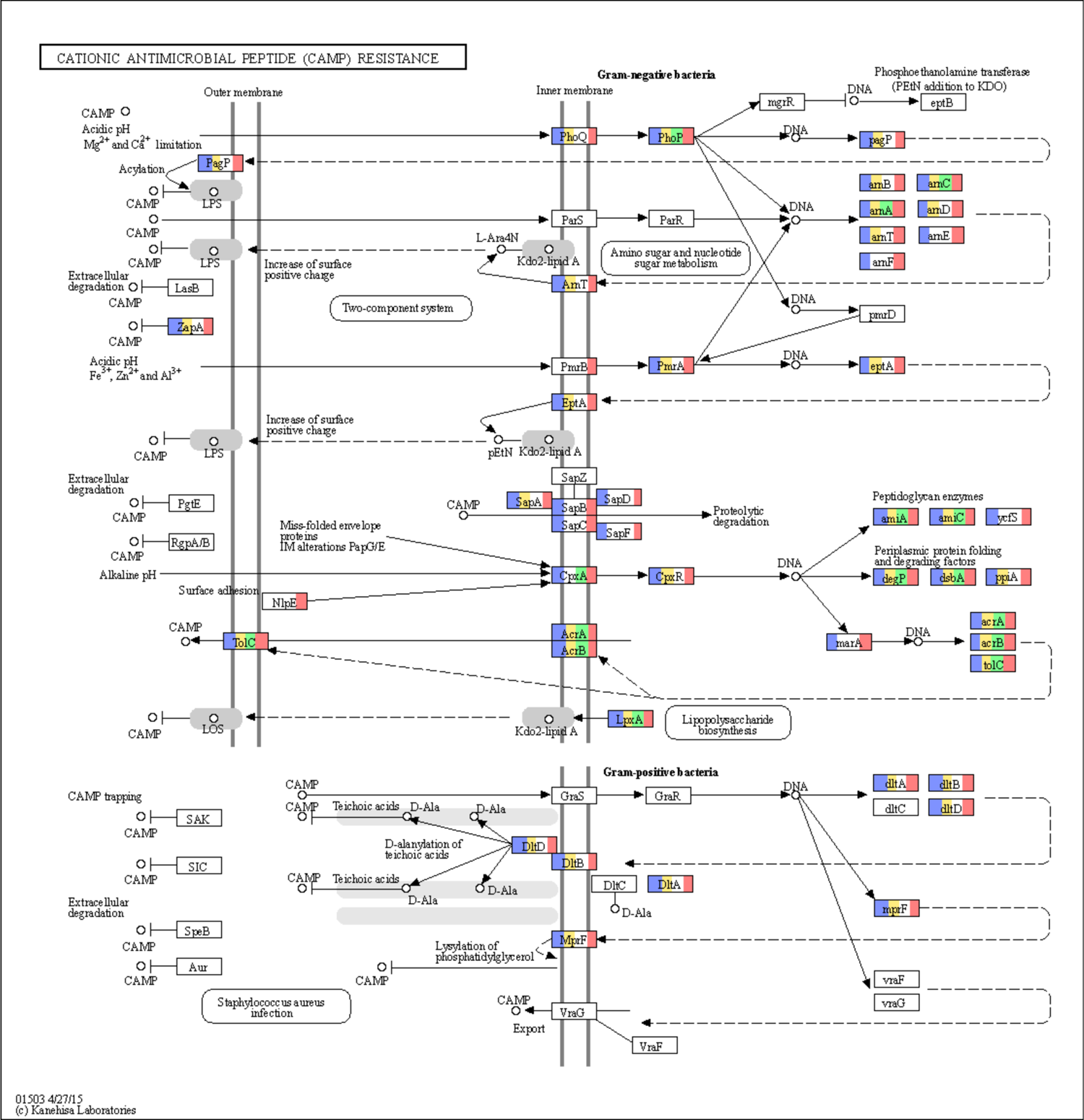
Cationic antimicrobial peptide (CAMP) resistance pathway based on KEGG orthology. Colors indicate the genes that were found to be present in a certain sample: influent iDNA (blue), influent exDNA (yellow), effluent iDNA (green), and effluent exDNA (red).

**Figure S8.**
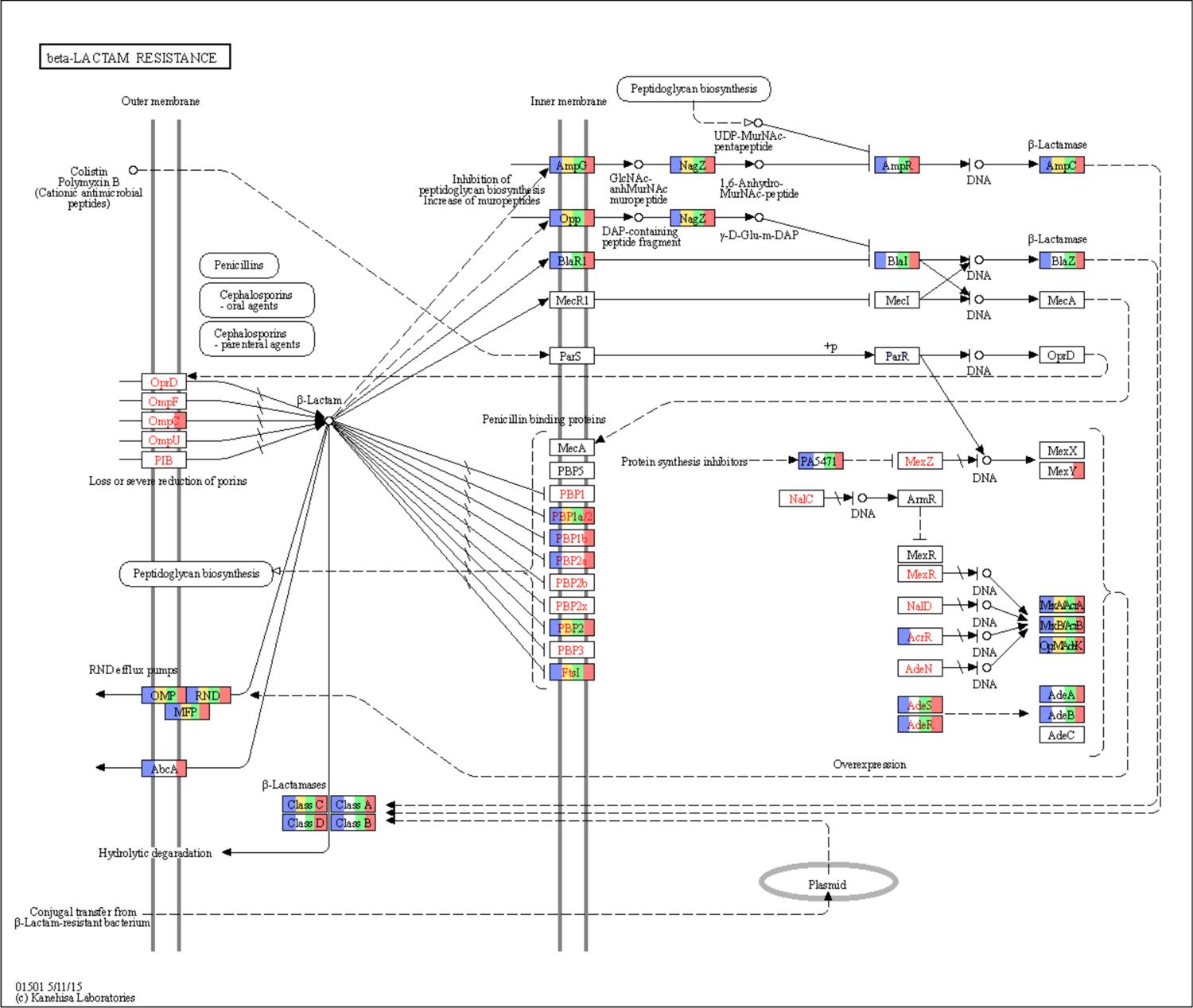
The beta-lactam resistance pathway based on KEGG orthology. Colors indicate the genes that were found to be present in a certain sample: influent iDNA (blue), influent exDNA (yellow), effluent iDNA (green), and effluent exDNA (red).

### Quantitative PCR supplementary results

**Figure S9.**
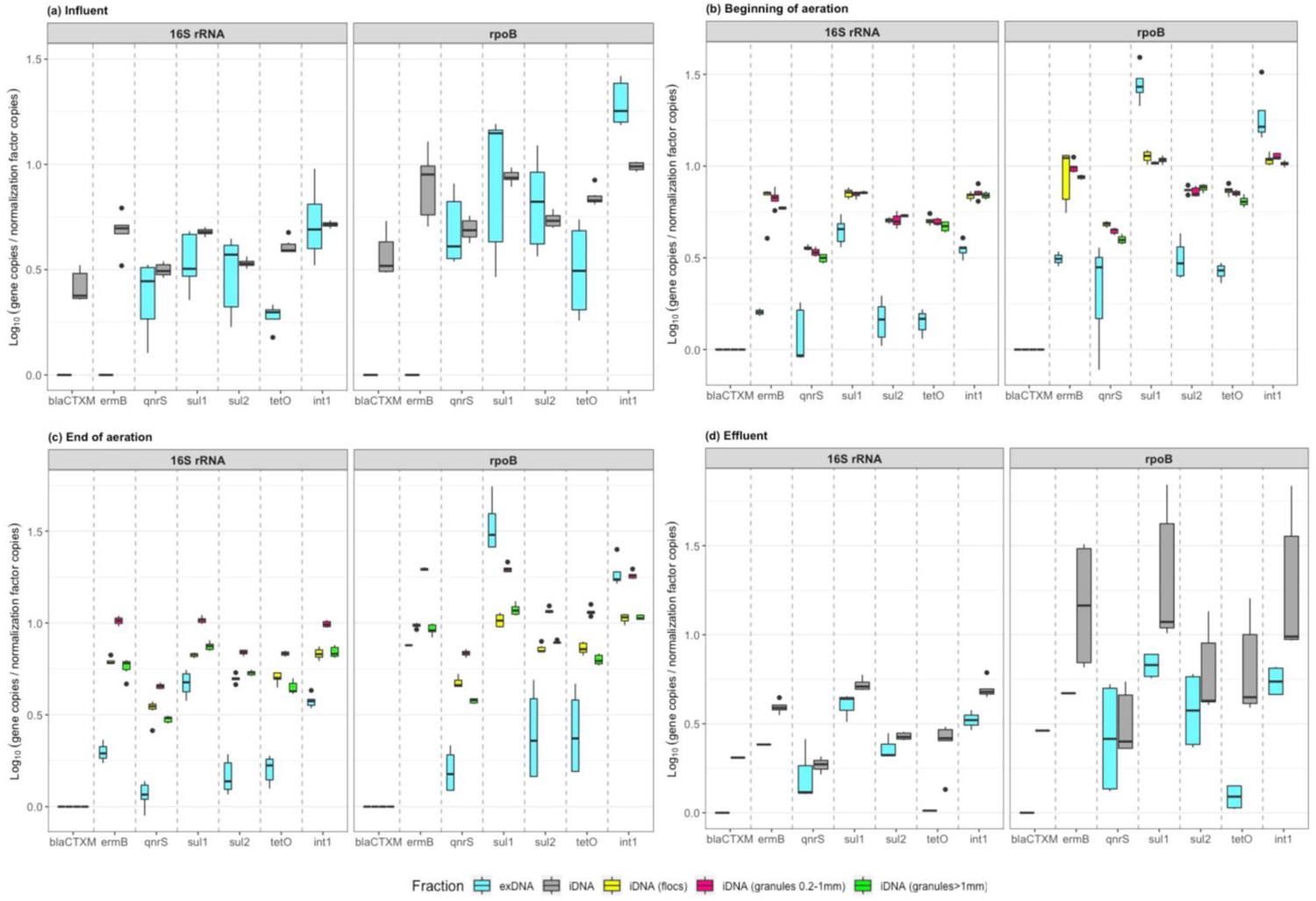
Relative abundance of gene copies using *16S rRNA* and *rpoB* as a proxy for the concentration of the bacteria. Genes are quantified in terms of log_10_ gene copies / log_10_ *16S rRNA* or *rpoB* gene copies. The four different graphs represent the four AGS WWT steps: (**a)** influent, **(b)** beginning of the aeration**, (c)** end of the aeration, and **(d)** effluent.

**Figure S10.**
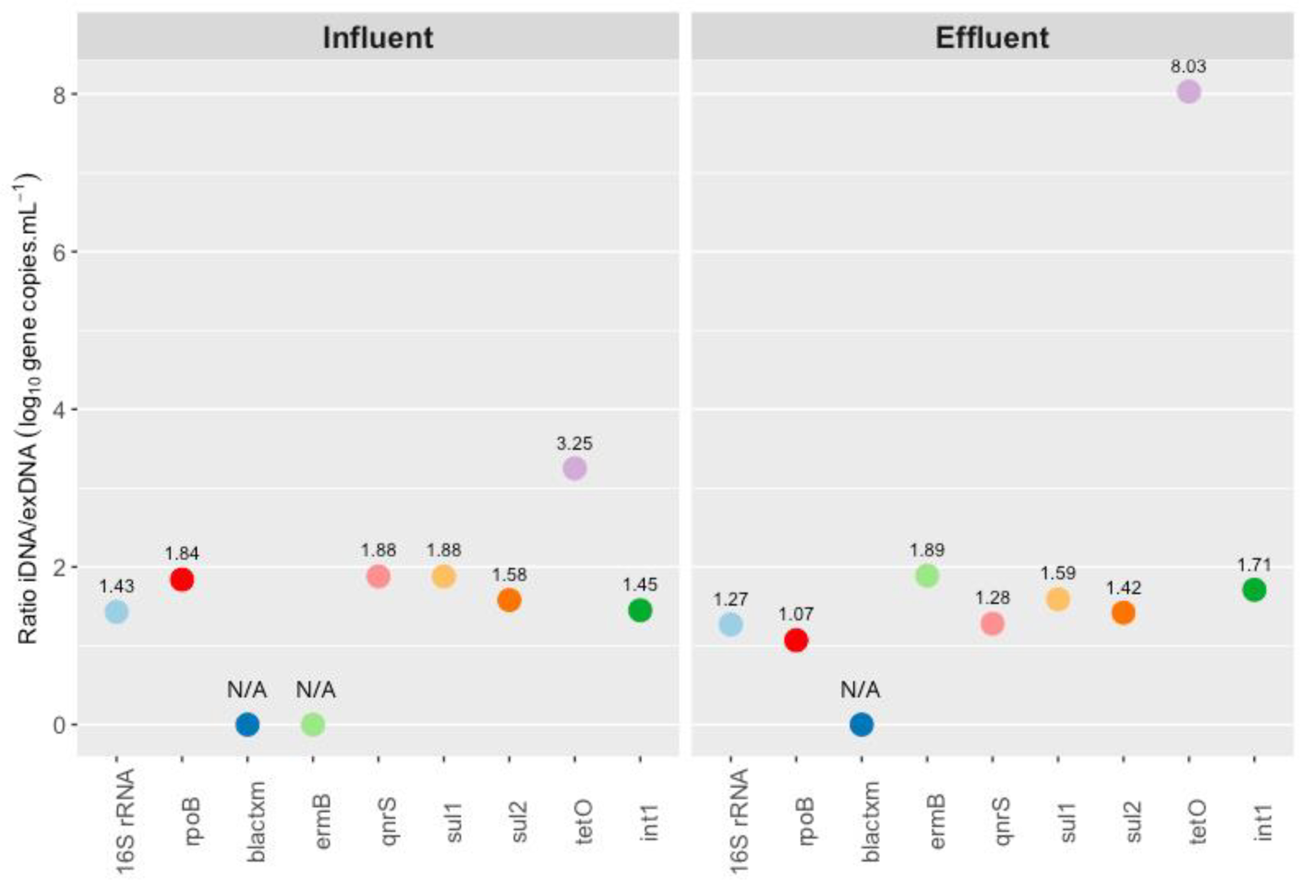
Ratio of intracellular over extracellular DNA concentration of the genes measured in log_10_ gene copies mL^-1^ for the influent and the effluent phase of the AGS WWT process. Ratios labelled with N/A indicate that the gene was not detected in at least one of the two DNA fractions.

**Figure S11.**
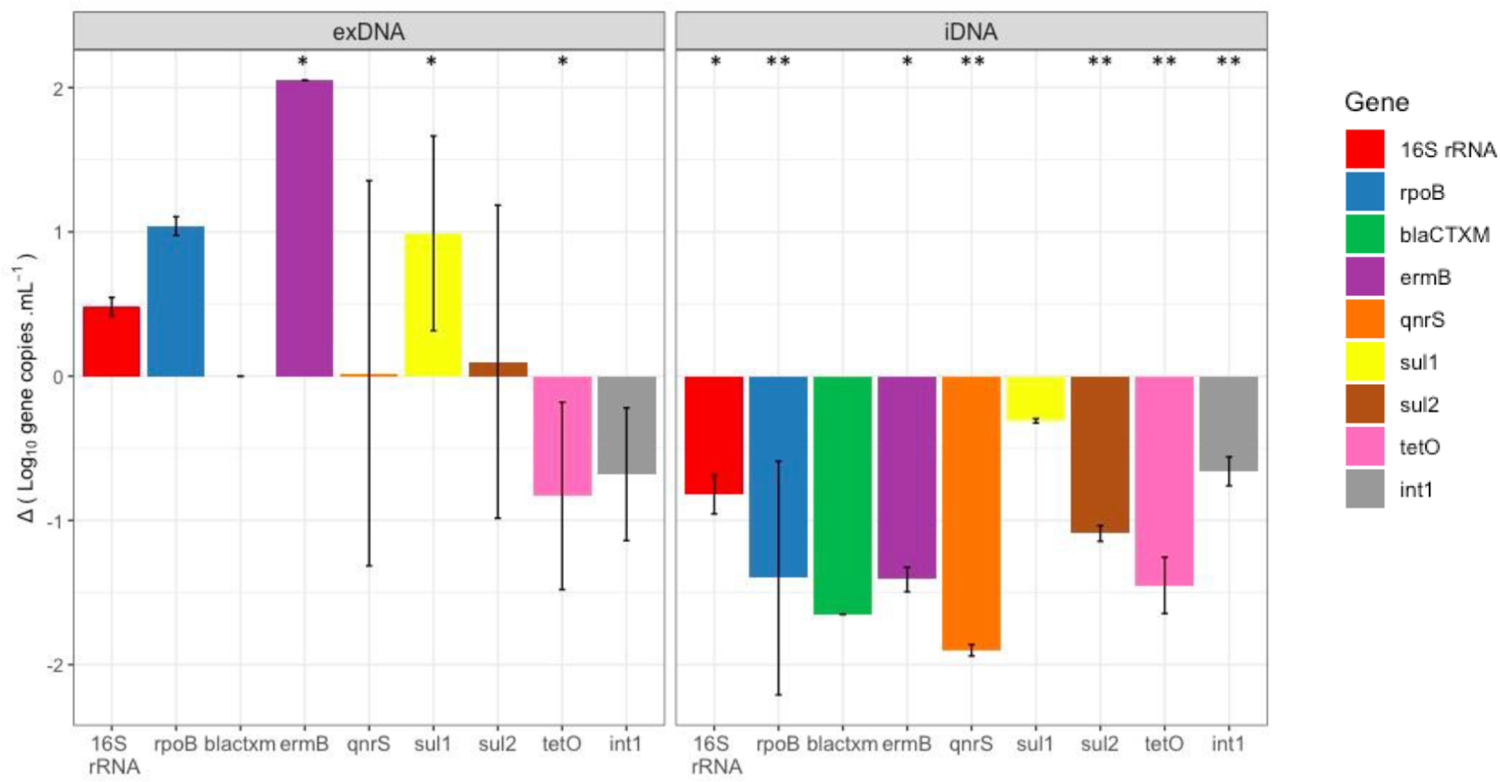
Effluent minus influent (Δ) in log_10_ gene copies mL^-1^ for the extracellular (left) and intracellular (right) DNA fraction. The significance of the data, as calculated by the non-parametric Wilcoxon test, is displayed as p<0.05 (* labels) and p<0.005 (** labels).

### ARGs and MGEs co-localization results

**Table S6.**
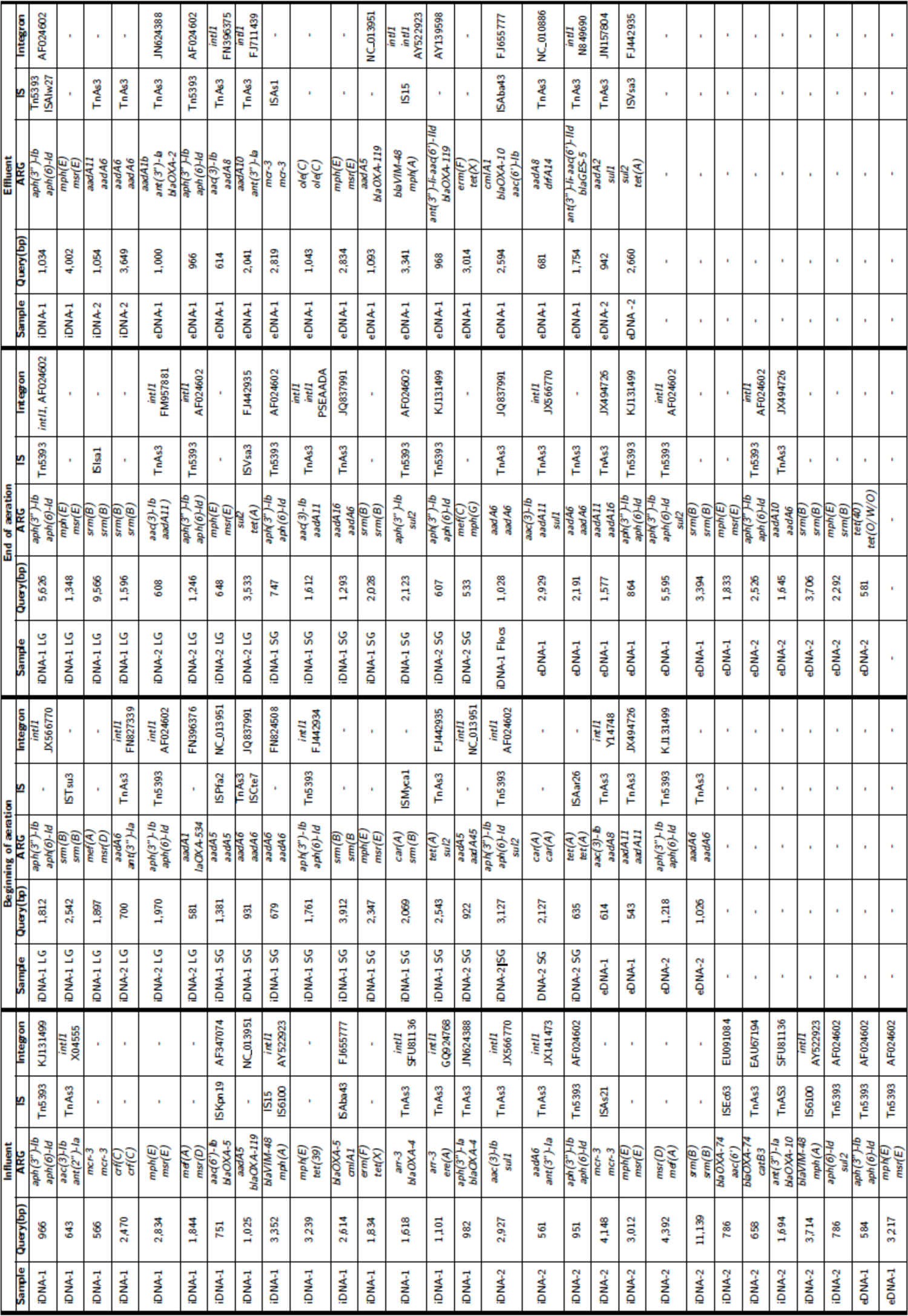
Identified contigs containing multiple ARGs. Some also contain MGEs. Annotation of contigs by alignment in ISfinder database is provided in the “Insertion sequence” column with corresponding ISfinder nomenclature. Annotation of contigs by alignment in INTEGRAL database is provided in the “Integron” column with corresponding NCBI accession number.

**Table S7.**
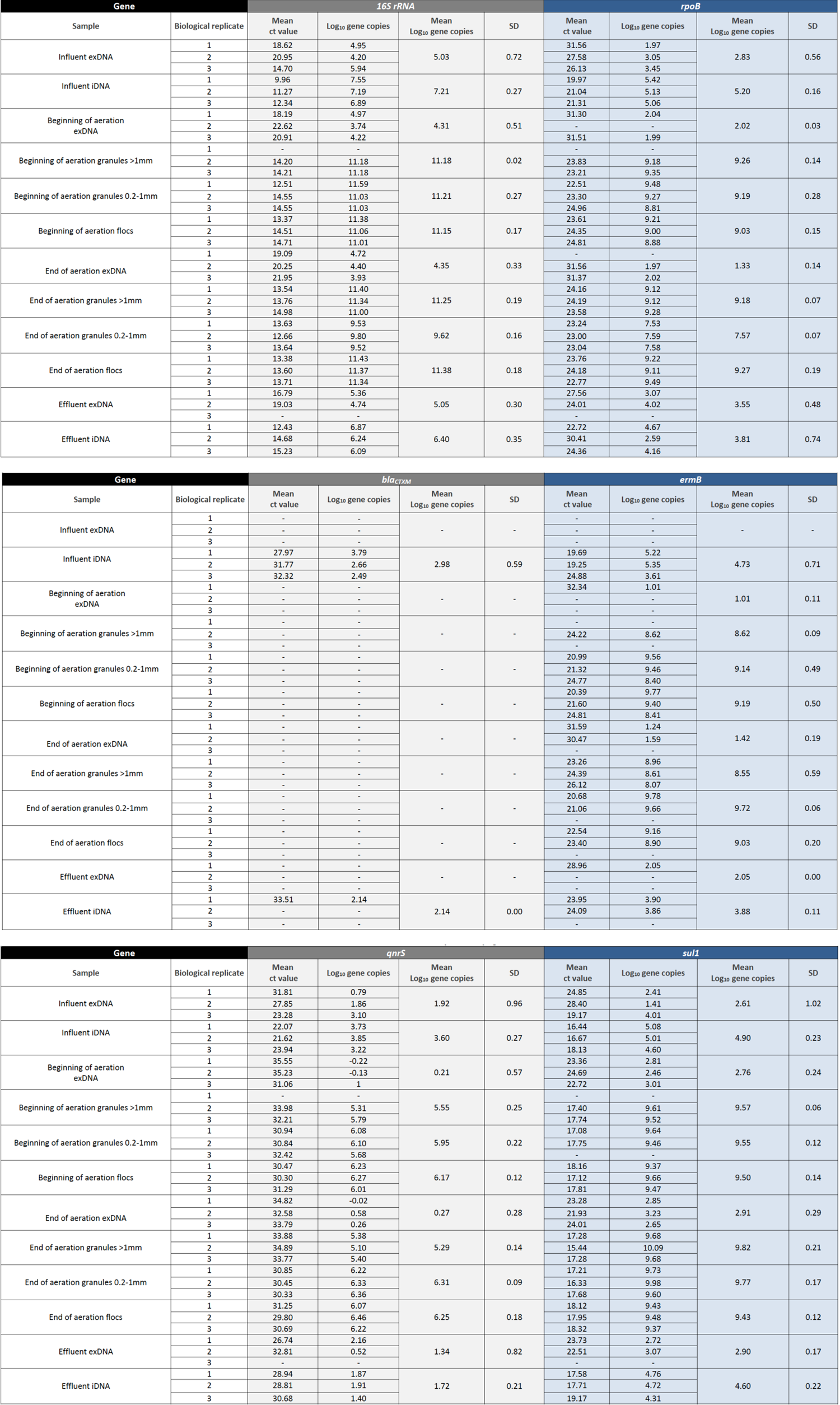
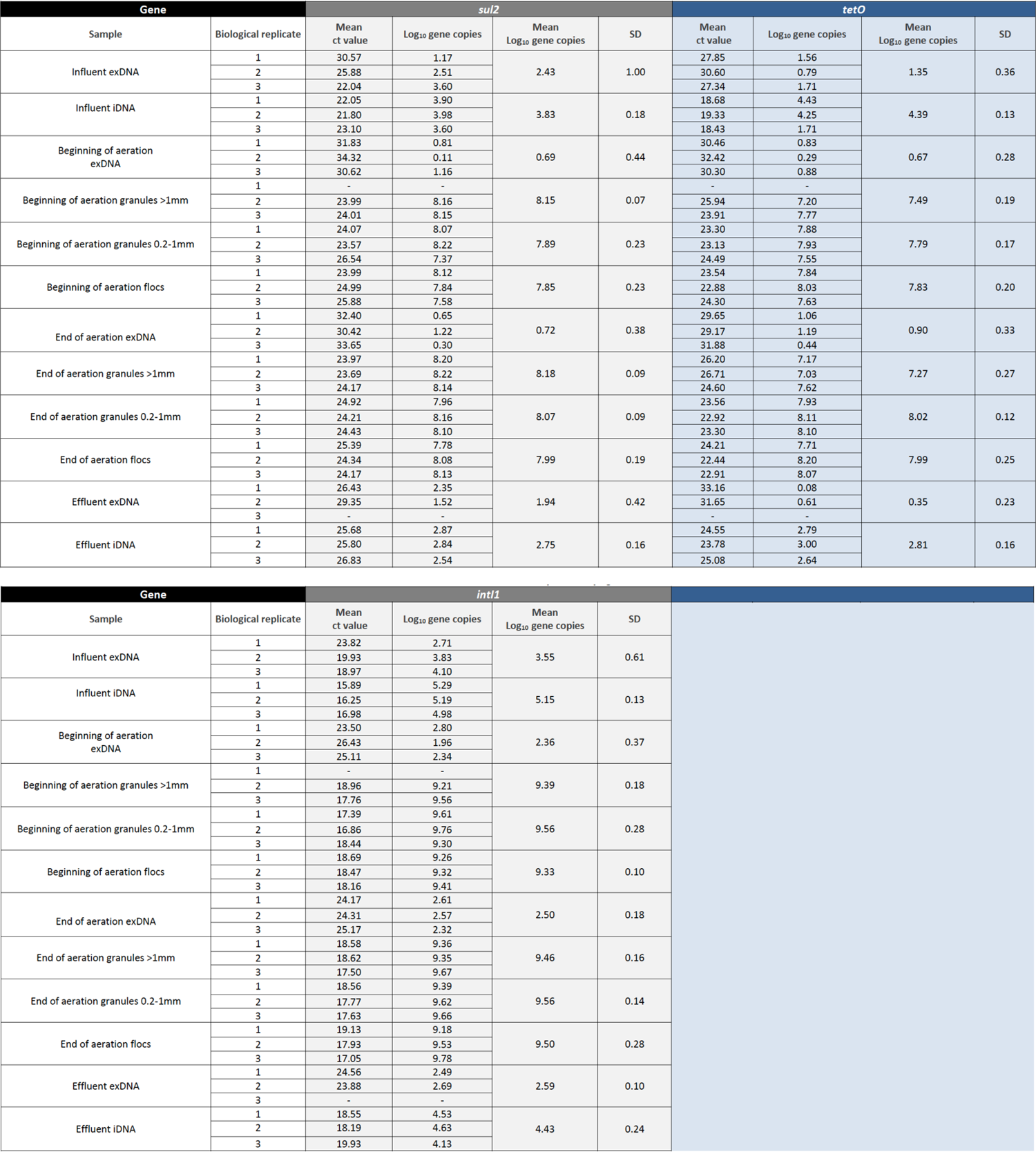
qPCR results for the selected panel of ARGs and MGE during the AGS WWT operation process.

**Table S8.**
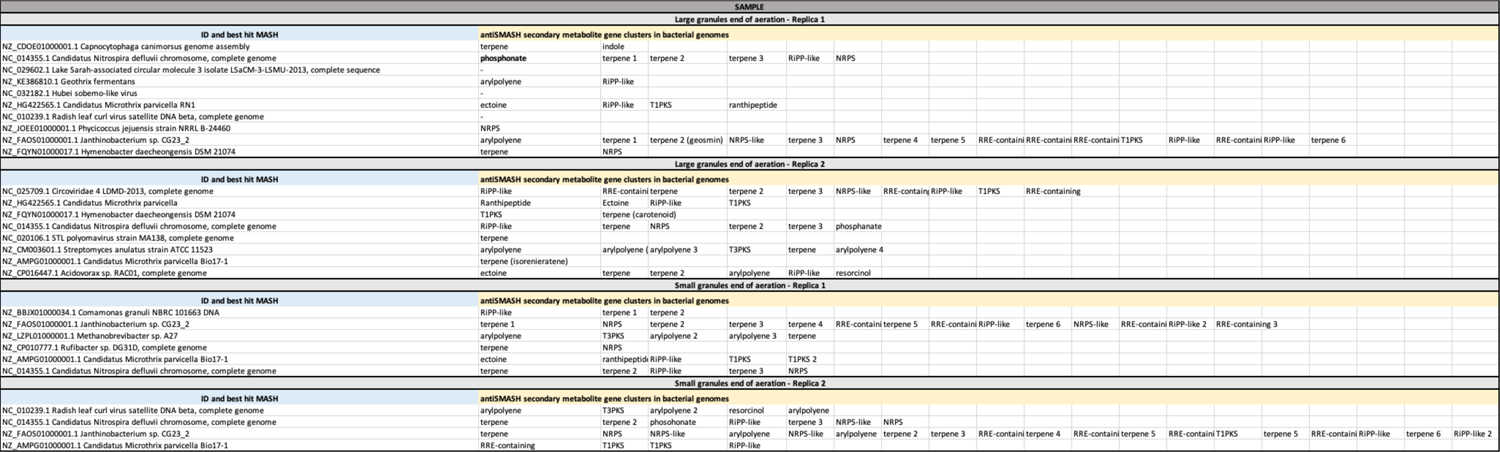
Secondary metabolite gene clusters identified from recovered metagenome-assembled genomes (MAGs) from small and large granules at the end of the aeration of the AGS WWT operation process in both bioreplicates. MAGs from floccular sludge could not be recovered.

